# Femtosecond laser microdissection isolating regenerating *C. elegans* neurons for single cell RNA sequencing

**DOI:** 10.1101/2021.01.21.427576

**Authors:** Peisen Zhao, Chris Martin, Ke-Yue Ma, Ning Jiang, Adela Ben-Yakar

## Abstract

Our understanding of nerve regeneration can be enhanced by delineating its underlying molecular activities at single neuron resolution in small model organisms such as *Caenorhabditis elegans*. Existing cell isolation techniques cannot isolate regenerating neurons from the nematode. We present femtosecond laser microdissection (fs-LM), a new single cell isolation method that dissects intact cells directly from living tissue by leveraging the micron-scale precision of fs-laser ablation. We show that fs-LM facilitated sensitive and specific gene expression profiling by single cell RNA-sequencing, while mitigating the stress related transcriptional artifacts induced by tissue dissociation. Single cell RNA-sequencing of fs-LM isolated regenerating *C. elegans* neurons revealed transcriptional program leading to successful regeneration in wild-type animals or regeneration failure in animals lacking DLK-1/p38 kinase. The ability of fs-LM to isolate specific neurons based on phenotype of interest allowed us to study the molecular basis of regeneration heterogeneity displayed by neurons of the same type. We identified gene modules whose expression patterns were correlated with axon regrowth rate at a single neuron level. Our results establish fs-LM as a highly specific single cell isolation method ideal for precision and phenotype-driven studies.

## Main

Spinal cord injuries result in permanent functional deficit as axons in the adult central nervous system fail to regenerate after trauma^1^. State-of-the-art interventions promote neuron survival and axon regrowth by engaging neuron-intrinsic and pro-regenerative pathways^2^. However, clinical outcome of such interventions are still poor: regrowth can only be stimulated in a small number of neurons, while a seemingly homogeneous neuron population can exhibit diverse or even opposite responses to the same intervention^3, 4^. A deeper understanding of nerve regeneration can be attained by dissecting the process at single neuron resolution in a well-defined nervous system. For such studies, *Caenorhabditis elegans* has been established as a valuable model organism in conjunction with femtosecond laser axotomy^5, 6^, which can reproducibly induce a variety of relevant regeneration phenotypes *in vivo*. The transparent body of *C. elegans* further allows analysis of such phenotypes, including axon regrowth rate, guidance, and fusion at a single neuron resolution *in vivo* with fluorescence microscopy. Past nerve regeneration studies in *C. elegans* have led to the discovery of many conserved nerve regeneration pathways^7–26^, notably the dual leucine zipper kinase (DLK-1/p38) signaling cascade, whose role in neural development, regeneration, and degeneration has been confirmed in numerous vertebrate and invertebrate species^7, 27–31^.

Nerve regeneration research in *C. elegans* has relied on costly and time-consuming mutant screens that test individual genes for function in axon regeneration (Supplementary Note 1). The rapidly evolving high-throughput sequencing methods offer an opportunity to accelerate progress. Nevertheless, isolating regenerating neurons from *C. elegans* has so far remained infeasible. As in other models, the prevalent cell isolation method for *C. elegans* entails fluorescence activated cell sorting preceded by lengthy and disruptive tissue dissociation (dissociation-FACS method)^32^. Several drawbacks, notably low yield of cells (< 0.5%)^33^, induction of transcriptional artifacts^34, 35^, and inability to isolate specific neurons based on regeneration phenotypes, make this method inapplicable (Supplementary Note 2). Other cell isolation methods like laser capture microdissection (LCM) and Patch-seq^36^ can bypass tissue dissociation, but may cause intolerable contamination and degradation of cellular content, especially considering the small size of *C. elegans* neurons (< 3 µm in diameter with < 0.1 pg total RNA).

Above challenges motivated us to develop femtosecond laser microdissection (fs-LM) to resect intact single cells directly from living tissue. We demonstrate that the micron-scale precision of fs-laser ablation allows us to isolate specific neurons from living *C. elegans* in their intact form. The cell yield of fs-LM was 32%, which represented orders of magnitude improvement over the dissociation-FACS method. Importantly, fs-LM mitigated the stress related transcriptional artifacts induced by the tissue dissociation protocol. We used fs-LM to isolate one specific type (posterior lateral microtubule, PLM) of *C. elegans* neuron, exhibiting successful or failed regeneration in with-type or DLK-1 knockout animals following fs-laser axotomy. Single cell RNA sequencing (scRNA-seq) of the isolated neurons revealed transcriptional program required for axon regeneration and its potential upstream regulators. To investigate the molecular basis of regeneration heterogeneity displayed by neurons of the same type, we also isolated single regenerating PLM neurons on the basis of axon regrowth rate. We identified genes whose expression pattern correlated with axon regrowth rate at a single neuron resolution. Weighted gene co-expression network analysis further revealed that accelerated axon regrowth requires elevated gene expression associated with functions like neuorogenesis, DNA repair, and RNA transcription, as well as reduced gene expression involved in immune response and RNA metabolism. Our results establish that by resecting intact cells from living tissue with fs-laser ablation, fs-LM achieves precise isolation of specific cells on the basis of phenotype of interest. Such capability uniquely positions fs-LM to facilitate precision and phenotype-driven studies for deciphering complexity of tissue. To show versatility of fs-LM beyond *C. elegans*, we also demonstrated isolation of single cortical neurons from acute mice brain slices.

## Results

### fs-LM isolates single intact touch receptor neurons from living *C. elegans*

Laser microdissection of single cells from living tissue is highly challenging as collateral damage from laser ablation can easily damage the cells being isolated. The challenge is even greater when isolating single neurons from *C. elegans* due to the small size of its neurons (< 3 µm in diameter). To meet this challenge, we leveraged the precision of ultrafast laser microsurgery, in which high peak intensity of tightly focused fs-laser pulses allows for sub-micron scale tissue ablation. Such precise ablation process can greatly mitigate both collateral and out-of-focus tissue damage, which our group has previously demonstrated with fs-laser axotomy in *C. elegans^5,6,37,38^*.

To isolate single neuron from *C. elegans*, fs-LM first resects the target neuron by ablating its surrounding tissue at a series of spots arranged in a 3D hemispherical shell encasing the neuron (Fig. 1a, b). The ablation spots are spaced 2 – 3 µm apart from each other and the target neuron. At each spot, a train of low-energy laser pulses is delivered to ablate a small volume (< 2 µm) of tissue (Supplementary Note 3). With such ablation pattern, we can effectively remove tissue in the immediate vicinity of the neuron while keeping its soma intact. To release the resected neuron, we ablate cuticle of the nematode at 5 µm from the neuron to create a 2 µm wide incision. The neuron is then promptly released through the incision driven by the internal pressure of *C. elegans* (Fig. 1a, b, Supplementary Fig. 1a-d). Each released neuron is visually inspected for adherent debris, which can be removed by laser-induced bubbles as needed. We then collect the released neuron with a micropipette and bath it in cleaning buffer twice to remove any remaining contaminant in the aspirate. Finally, the cleaned neuron is deposited into lysis buffer and frozen on dry ice.

**Fig. 1.**
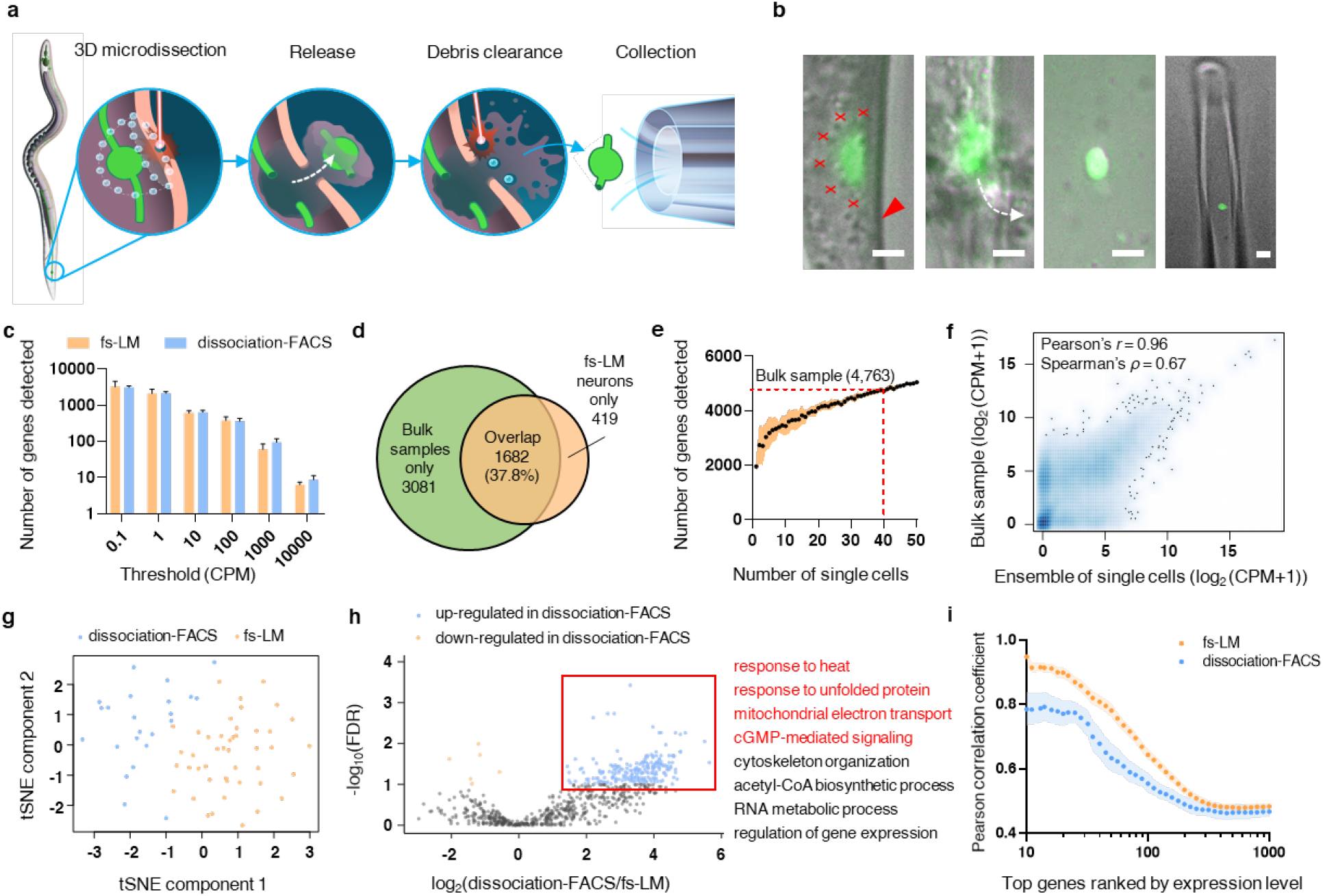
fs-LM isolates intact neurons with reduced stress related transcriptional artifacts for sensitive and specific gene expression profiling by scRNA-seq. **a**, Schematic description of the fs-LM method. **b**, Microscope images of fs-LM during isolation of a GFP-labeled PLM neuron in the order corresponding to the schematic in (a). Red crosses: locations of laser ablation spots at the current focal plane. Spots on all focal planes form a hemispherical shell encasing the neuron. Red arrowhead: location of cuticle ablation. White dashed line: trajectory of neuron release. Scale bar: 2 µm. **c**, Number of genes detected at various thresholds in single neurons isolated by fs-LM or dissociation-FACS method. Error bars represent 95% confidence interval. **d,** Overlap between genes identified in fs-LM isolated single neurons and bulk sample (> 5,000 FACS neurons). **e,** Number of genes detected from pooled scRNA-seq reads from different numbers of fs-LM isolated neurons. Shaded area represents 95% confidence interval. **f,** Correlation of gene expressions between ensemble of 40 fs-LM isolated neurons and bulk sample. **g**, t-SNE visualization of gene expression in neurons isolated by fs-LM and dissociation-FACS methods. **h**, Volcano plot showing differential gene expression with orange and blue data points highlighting the significantly down-regulated and up-regulated genes, respectively, in dissociation-FACS isolated neurons. Statistical significance was calculated as Benjamini-Hochberg false discovery rate (FDR). In this test for differential gene expression, a threshold of 10% FDR (FDR < 0.1) was used. The list shows representative gene ontology terms (FDR < 0.001) associated with the up-regulated genes (red window), including key functions related to stress response (highlighted in red). **i,** Correlation among gene expression profiles of single neurons isolated by fs-LM or dissociation-FACS methods, showing a significantly stronger correlation among fs-LM isolated neurons (two-way ANOVA test, *P* < 0.0001). Shaded areas represent 95% confidence interval.

To demonstrate feasibility of fs-LM, we isolated posterior lateral microtubule (PLM) neurons from living larval stage (L4) *C. elegans*. A typical fs-LM took 4 – 5 minutes per neuron with a yield of 32% (*n* = 384). Such high yield represented orders of magnitude improvement over the dissociation-FACS method (< 0.5%, Supplementary Note 2). We used cytoplasmic GFP as a convenient indicator of neuron structural integrity and only collected neurons that retained their GFP signal level after being released (Fig. 1b, Supplementary Fig. 1a-d). In case of neuronal damage, which primarily occurred as neurons became torn or fragmented during the release process due to incomplete resection, GFP signal of the neuron dropped drastically (Supplementary Fig. 1e-l). During dissection, we avoided cutting the axons close to the neuron, which would cause significant loss of cellular content. Instead, we severed the axon at a safe distance of 20 µm away from the soma, which promptly sealed after being cut as we previously showed^6^.

We constructed scRNA-seq libraries from 94.0% of the fs-LM isolated PLM neurons (*n* = 123) that passed cDNA quality check. The failed samples were mostly a result of neuron lysis while still in the micropipette and were excluded (Supplementary Fig. 2). In addition, we collected both pooled and single touch receptor neurons with the dissociation-FACS method according to established protocol (Supplementary Fig. 3)^32^. Sufficient sequencing depth was attained for 90% of the sequenced single neurons (Supplementary Fig. 3a-c). We detected an average of 2,261 ± 132 genes by a threshold of 1 count per million (CPM) reads in each fs-LM isolated neuron, accounting for 37.8% of all expressed genes in touch receptor neurons, comparable to the dissociation-FACS method^39, 40^ (Fig. 1c,d).

Tissue enrichment analysis of the genes detected in fs-LM isolated neurons identified touch receptor neuron as the top enriched term (adjusted *P* < 10^−20^, Supplementary Fig. 4d, e). We also identified known marker genes specific to PLM neurons, such as *mec-17, mec-12, mec-7, mec-10*, and *mec-18*, in fs-LM isolated neurons (Supplementary Fig. 4f). As we expect fs-LM to be applied for gene expression profiling of rare cell types that cannot be acquired in a high-throughput manner, we tested whether fs-LM isolated cells can recapitulate bulk samples. We found that by pooling scRNA-seq reads from 40 fs-LM isolated neurons (Fig. 1e), the number of genes detected reached that of bulk sample. Furthermore, gene expression profiles exhibited good correlation between the ensemble of single neurons and the bulk sample (Pearson correlation coefficient = 0.96, Fig. 1f). Our results confirm that as a new single cell isolation method, fs-LM yields intact single neurons ideal for sensitive and specific gene expression profiling using scRNA-seq.

### fs-LM mitigates transcriptional artifacts and improves reproducibility of scRNA-seq

Past studies have found induction of stress response genes among cells isolated by the dissociation-FACS method. Such transcriptional artifacts are a result of the tissue dissociation protocol, which involves a prolonged enzymatic digestion at elevated temperatures, and can severely affect interpretation of data^34, 35^. fs-LM can mitigate such artifacts by minimizing stress to the target cell thanks to the high precision of fs-laser ablation. In addition, as early transcriptional response peaks at 25 minutes following stimuli^41, 42^, the relatively short fs-LM protocol (< 5 minutes) can be completed before the artifacts become significant. We compared gene expression in touch receptor neurons isolated by fs-LM and dissociation-FACS methods. As expected, neurons isolated by different methods formed two clusters following t-SNE analysis (Fig. 1g), indicating that transcriptome of the isolated neurons were indeed affected by isolation methods.

Next, we investigated the differential gene expression between neurons isolated by these two methods (Fig. 1h). Using gene expression in fs-LM isolated neurons as reference, we found 219 up-regulated genes and 6 down-regulated genes among dissociation-FACS isolated neurons (FDR < 0.1, Supplementary Table 2). Gene Ontology (GO) analysis of the up-regulated genes identified stress related terms such as response to heat (FDR = 8.3 x 10^−7^), cellular response to unfolded protein (FDR = 3.5 x 10^−5^), and mitochondrial electron transport chain (FDR = 1.2 × 10^−4^). Among the top up-regulated genes, we identified genes that were also highly up-regulated during heat shock response in a manner dependent on heat shock factor 1^43^, such as Y43F8B.2, which is involved in innate immune response, and genes encoding heat shock proteins (e.g. F44E5.5, F44E5.4, *hsp-16.1, hsp-16.41*, and *hsp-16.48*, Supplementary Fig. 5b, d). The up-regulated genes also included mtDNA-encoded *nduo-2, nduo-3, nduo-5*, and *nduo-6*, which have NADH dehydrogenase activity and were up-regulated in mitochondrial unfolded protein response^44^. Meanwhile, the down-regulated genes included *mec-7, mec-12*, and *mec-17*. The MEC-12 α-tubulin and the MEC-7 β-tubulin compose the 15 protofilament microtubules (MTs) in *C. elegans* touch receptor neurons and are critical for touch sensitivity. MEC-17 acetyltransferase functions in acetylation of K40 on MEC-12, which is required for MT stability and axonal transport^45^. These genes were found to be down-regulated when the MTs were disrupted by colchicine^28, 46^. The observed pattern of gene regulation among dissociation-FACS isolated neurons exhibited significant alignment with the pattern previously profiled in heat shock response^43^ (Supplementary Fig. 5). The results above indicate that compared to fs-LM isolated neurons, dissociation-FACS isolated neurons display stronger induction of stress response, thus suggesting that fs-LM can alleviate stress related transcriptional artifacts among isolated cells.

We further examined if fs-LM can improve reproducibility of scRNA-seq compared to the dissociation-FACS method. Our results indicated that fs-LM isolated neurons showed significantly higher correlation in expression levels (two-way ANOVA test, *P* < 0.0001), especially among the highly expressed genes (Fig. 1i), which was expected since fs-LM can avoid technical variability introduced by stress related transcriptional artifacts. Moreover, fs-LM can isolate one specific neuron out of the six touch receptor neurons in each animal, while the dissociation-FACS method can only isolate all six neurons simultaneously as they are all GFP-labeled by the same promoter. Together, the results above indicate that fs-LM can mitigate transcriptional artifacts induced by the dissociation-FACS method and improve reproducibility of scRNA-seq.

### fs-LM facilitates profiling of transcriptional dynamics underlying axon regeneration in *C. elegans*

With fs-LM, we were able to isolate regenerating *C. elegans* neurons, which has not been possible with existing methods. In wild-type animals, axons of axotomized PLM neurons initiate regrowth from their proximal stump at 4 – 6 hours post axotomy, leading to distinct regeneration outcomes (Fig. 2a, Supplementary Fig. 6). To avoid confounding gene expression induced by regeneration with the immediate injury response to axotomy^47^, we isolated regenerating neurons at 24 hours post axotomy. Only neurons that displayed no reconnection to the distal fragment were collected, since potential axonal fusion can trigger restorative processes that alter the neurons’ transcriptional state^48^. We also isolated axotomized PLM neurons from *dlk-1 (0)* mutants at 24 hours post axotomy, which failed to initiate regrowth by forming a growth cone. As control, we isolated uninjured PLM neurons from both wild-type and *dlk-1 (0)* animals that underwent mock axotomy.

**Fig. 2.**
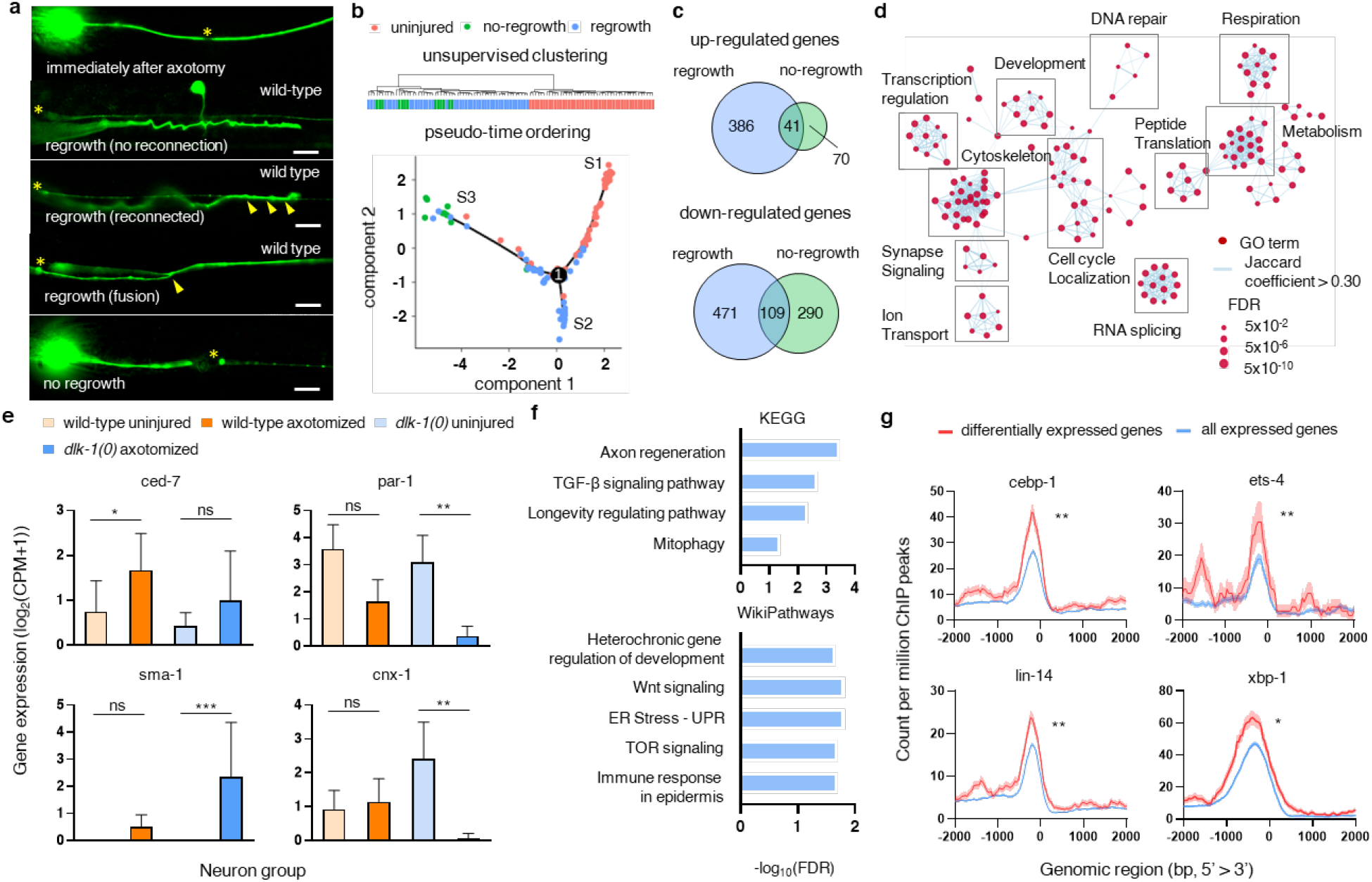
fs-LM facilitates profiling of transcriptional program underlying nerve regeneration in *C. elegans*. **a**, Microscope images of axotomized PLM neurons immediately after axotomy and 24 hours post axotomy, with regrowth & no reconnection, regrowth & reconnection, and regrowth & fusion phenotypes in wild-type animals and no-regrowth phenotype in *dlk-1 (0)* mutants. Yellow asterisks indicate sites of axotomy, and the yellow arrowheads indicate sites of reconnection to the distal fragment. Scale bar: 10 µm. **b**, Unsupervised hierarchical clustering (top) and pseudotemporal ordering (bottom) of uninjured, axotomized wild-type (regrowth), and axotomized *dlk-1 (0)* (no-regrowth) neurons based on 1,000 genes with highest variance in expression levels. The uninjured neurons from both wild-type and *dlk-1 (0)* animals are represented by red since they showed few DEGs (Supplementary Fig. 7a). **c,** Overlap between DEGs in successful or failed regeneration. Blue circles (regrowth) represent DEGs between axotomized and uninjured neurons from wild-type animals. Green circles (no-regrowth) represent DEGs between axotomized and uninjured neurons from *dlk-1 (0)* animals. **d**, Network visualization of GO analysis results. Red nodes represent enriched GO terms (FDR < 0.01). Network edges represent pairwise semantic overlap among the GO terms (Jaccard coefficient > 0.3). **e**, Representative regeneration regulators with distinct expression patterns across different neuron groups. Asterisks: Statistical significance (*: 0.01 < FDR < 0.05; **: 0.001 < FDR < 0.01; ***: FDR < 0.001; ns: not significant). Error bars represent 95% confidence interval. **f**, GO analysis of transcription factors with significant binding affinity to the DEGs. **g**, Metagene binding profiles of representative transcription factors centered at the transcription start site. Red traces represent density of chromatin immunoprecipitation sequencing (ChIP) peaks mapped to the DEGs. Blue traces represent density ofChIP peaks mapped to all genes expressed in touch receptor neurons. Asterisks: Statistical significance (*: 0.01 < *P* < 0.05; **: 0.001 < *P* < 0.01; ***: *P* <0.001; ns: not significant). Shaded area represents 95% confidence interval.

To understand the transcriptional dynamics leading to successful regeneration in wild-type animals and regeneration failure in *dlk-1 (0)* animals, we performed scRNA-seq on the collected neurons. Unsupervised hierarchical clustering showed that the uninjured and axotomized neurons formed two clusters independent of their genetic background, indicating that the observed transcriptional activities were dominated by axon regrowth instead of DLK-1 knockdown (Fig. 2b). Further pseudotemporal ordering by Monocle2^49^ positioned the neurons into 3 branches according to their regeneration states (Fig. 2b). S1 branch comprised mostly uninjured neurons from both wild-type and *dlk-1 (0)* animals, S2 branch comprised mostly regrowing neurons from wild-type animals, and S3 branch included most of the non-regrowing neurons from *dlk-1 (0)* animals. The single branching point suggested a transition from an uninjured state into two distinct transcriptional states of successful or failed regeneration.

We profiled the differentially expressed genes (DEGs) between axotomized and uninjured neurons from each strain (Supplementary Fig. 7). Between the regrowing and uninjured neurons from wild-type animals, we identified 1,007 DEGs (fold change > 2, FDR < 0.05), including 427 up-regulated and 580 down-regulated genes (Fig. 2c, Supplementary Table 3). Between the non-regrowing and uninjured neurons from *dlk-1 (0)* animals, we identified 510 DEGs, most of which were down-regulated. The 150 overlapping DEGs between these 2 sets exhibited significant correlation in fold change of expression levels (Pearson correlation coefficient = 0.94, *P* = 2 × 10^−16^), suggesting that these DEGs were likely induced by stress response or loss of neuronal function instead of axon regeneration (Supplementary Fig. 8a). The non-overlapping DEGs were likely involved in axon regeneration, with their selective induction leading to successful or failed axon regeneration. GO analysis followed by network visualization of these DEGs indeed revealed major network hubs associated with relevant terms such as cytoskeleton, synapse signaling, and development (Fig. 2d), which were also identified in regenerating mammalian neurons^50^.

### fs-LM facilitates discovery of genetic programs driving axon regrowth in *C. elegans*

To further understand the regulatory mechanisms involved in axon regeneration, we first investigated the links between the identified DEGs and the known regeneration mechanisms in *C. elegans*. We compiled a list of over 300 known regeneration regulators in *C. elegans* (Supplementary Table 1) and identified 48 regulator genes overlapping with the DEGs (*P* = 0.0015, Chi-squared test with Yates correction; Supplementary Table 4), including 39 required regulators and 9 inhibitory regulators for axon regeneration. Interestingly, majority the required regulators were up-regulated in regrowing neurons (30/39, *P* < 0.0001, Chi-square test) and down-regulated in non-regrowing neurons (23/39, *P* = 0.0024, Chi-square test).

The differentially expressed regeneration regulators exhibited a variety of previously unknown expression patterns following axotomy, which added new insights to their regulatory mechanism (Fig. 2e, Supplementary Fig. 8b). For example, *ced-7* encodes an ortholog of human ABC transporter and is required for axon regeneration. Its elevated expression following axotomy can promote axon regeneration by enhancing the recognition of axonal injury by exposing phosphatidylserine^21, 23^. Among the inhibitory regulators, we found PAR-1/MARK (MT affinity-regulating kinase). Down-regulation of *par-1* in axotomized neurons suggested adaptation of MT dynamics to axon injury, likely through localization of the EFA-6 ARF guanyl-nucleotide exchange factor^51^. Importantly, some regulators showed disparate expression patterns in wild-type and *dlk-1 (0)* animals. For example, *sma-1* encodes a β_H_-spectrin required for axon regeneration^52^. Selective up-regulation of *sma-1* in non-regrowing neurons is reminiscent of its role in constraining dendrite overgrowth post-embryonically^53^. *cnx-1* encodes an ortholog of calnexin, whose selective down-regulation in non-regrowing neurons suggests that abnormality in calcium release from internal storage likely contribute to the regeneration failure in *dlk-1 (0)* animals^8, 54^.

To explore the upstream regulators of the identified DEGs, we screened for transcriptional factors with significant binding affinities to the DEGs utilizing the chromatin immunoprecipitation sequencing (ChIP-seq) data from modENCODE and modERN repositories^55,56^ (Supplementary Fig. 9). We identified 38 transcription factors (TFs, *P* < 0.01) among the 204 screened *C. elegans* transcriptional factors (Supplementary Table 5). GO analysis for Kyoto Encyclopedia of Genes and Genomes (KEGG) pathways listed axon regeneration (*ets-4, cebp-1*, and *hif-1*) and TGF-β signaling pathway (*lin-35* and *sma-3*) as top enriched pathways.

The identified TFs included 6 known regeneration regulators (*cebp-1, ets-4, zip-4, unc-86, hif-1*, and *xbp-1*). Upon axon injury, CEBP-1 and ETS-4 function downstream to DLK-1/p38 and cAMP pathway, respectively, to form a complex which induces expression of *svh-2* gene^29^. Stimulation of SVH-2 receptor tyrosine kinase promotes axon regeneration through activation of the JNK pathway^31, 57^. Our data indicated an up-regulation of *svh-2* gene in both wild-type and *dlk-1 (0)* animals following axotomy, although the change was not statistically significant (data not shown). XBP-1 is a critical part of the endoplasmic reticulum unfolded protein response^58^. Recently, it is reported that a non-coding RNA produced from the cleavage of *xbp-1* mRNA is essential for axon regeneration in *C. elegans^59^*. The DEGs we identified likely contained downstream effectors of these regulator genes. Apart from the known regulators, we also identified TFs with potential novel regulatory roles in axon regeneration. For example, LIN-14 controls the positioning and embedment of axon in epidermis^60^. Adult *C. elegans* lacking *lin-14* exhibit axonal degeneration and guidance errors in both sensory and motor neurons. Our data suggests a role of *lin-14* in regulating transcriptional response to axon injury. The results above demonstrate that fs-LM can facilitate discovery of new insights on the regulatory mechanisms of nerve regeneration in *C. elegans*.

### fs-LM facilitates study of regeneration heterogeneity displayed by neurons of the same type

In both *C. elegans* and mammalian models, neurons display heterogeneity in regeneration capacity depending on neuronal type, age, preconditioning lesion, or injury condition^61^. Prior investigation into the molecular basis of such heterogeneity have largely focused on differential gene expression displayed by different neuronal types with varying regeneration capacities^62–64^. Nevertheless, significant regeneration heterogeneity also manifests among neurons of the same type^65, 66^. In *C. elegans*, the PLM neurons display regeneration heterogeneity hallmarked by a variable axon regrowth length at 24 hour post axotomy (Fig. 3a), even when experimental parameters such as animal age and injury conditions can be tightly controlled during fs-laser axotomy. Such regeneration heterogeneity is likely induced by random fluctuations in the intrinsic genetic machinery of axon regeneration. For example, prior work has shown that amplitude of calcium influx and release from internal storage following axon injury can modulate axon regrowth rate^8^. Unfortunately, despite its clinical importance as potential therapeutic target, this aspect of nerve regeneration has remained understudied due to a lack of method.

**Fig. 3.**
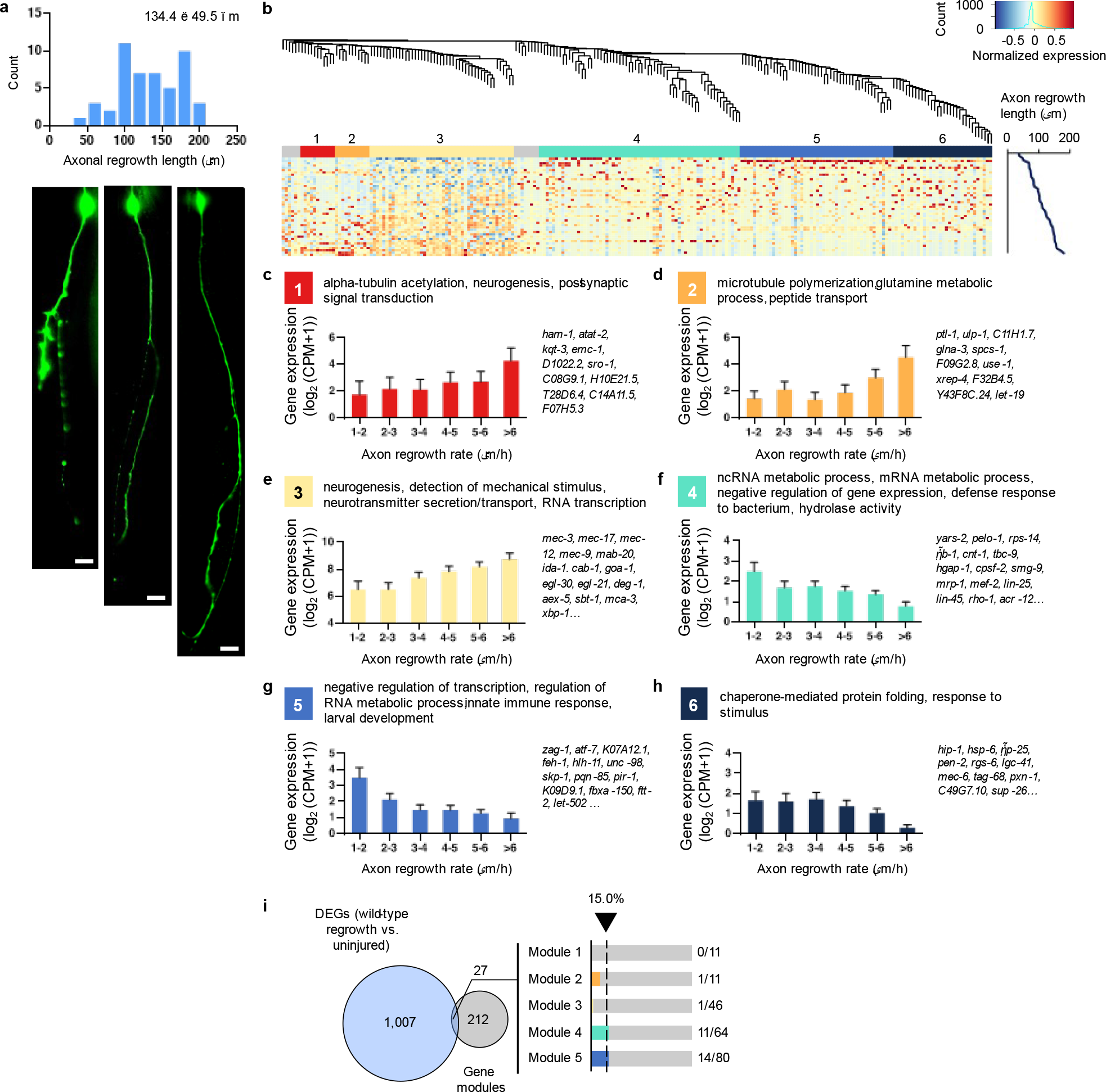
fs-LM facilitates study of regeneration heterogeneity displayed neurons of the same type. **a**, Histogram of axon regrowth lengths at 24 hours post axotomy (n=46) and microscope images of regenerating PLM neurons with different axon regrowth lengths. Scale bar: 10 µm. **b**, Weighted Gene Co-expression Network Analysis (WGCNA) of 226 candidate genes whose gene expression profile displayed significant correlation with axon regrowth rate. The dendrogram shows hierarchical clustering of the candidate genes into 6 co-expression modules, as labeled by the color band underneath. The heatmap shows expression levels of candidate genes across single regenerating neurons after min-max normalization. The rows represent single regenerating neurons, and are ordered according to the neurons’ axon regrowth rate, as indicated by the line chart on the right. **c-h**, GO terms associated with each of the 6 modules, bar plot showing the expression levels of all member genes corresponding to different axon regrowth rates (error bars: 95% confidence interval), and representative member genes within each module. **i**, Overlap between DEGs (regrowing versus uninjured wild-type neurons) and genes of the identified modules. Bar plots show the distribution of overlapping genes among the 6 modules and the percentage of genes within each module that overlaps with the DEGs.

With fs-LM, we were able to isolate regenerating PLM neurons on the basis of axon regrowth length at 24 hours post axotomy. We combined parametric and non-parametric tests to identify 226 candidate genes whose expression patterns significantly correlated with axon regrowth rate (Supplementary Fig. 10), with majority of them exhibiting a negative correlation (144/226, Supplementary Table 6). We further analyzed the co-expression patterns among the identified genes using weighted gene co-expression analysis (WGCNA)^67^. We identified 6 gene modules with distinct expression patterns and biological functions (Fig. 3b). GO analysis of each module revealed that accelerated axon regrowth is associated with elevated expressions of genes with functions related to neural development (Fig. 3c-e). Located in modules 1 – 3, these genes included microtubule related genes (*atat-2, mec-17, mec-12, mec-7, elp-1*, and *ptl-1*), ion channels and transporters (*kqt-3, mca-3*, and *deg-1*), synaptic transmission related genes (*snt-6, snt-4, cab-1, egl-21*, and *aex-5*), receptors (*emc-1, sro-1*, and *ida-1*), signaling molecules (*mab-20, egl-30, ndk-1, sbt-1, goa-1*, and *let-19*), and regulatory TFs (*ham-1* and *mec-3*). In addition, genes related to mechanosensory function (*mec-2, mec-3, mec-7, mec-9, mec-12*, and *mec-18*), DNA repair (*ulp-2* and *ulp-1*), and regulation of transcription (*mec-3, epc-1*, and *xbp-1*) were also up-regulated in fast regrowing neurons. Compared to module 1, module 2 displayed lower expression at regrowth rate < 5 µm/h, while module 3 showed higher overall expression yet smaller change with increasing axon regrowth rate.

In contrast, genes in modules 4 – 6 displayed reduced expression levels in fast regrowing neurons (Fig. 3f-h), including those related to negative regulation of transcription (*pelo-1, mrp-1, mef-2, zag-1*), ncRNA/mRNA metabolic process (*yars-2, pelo-1, fib-1, cpsf-2, smg-9* etc.), defense response (*lin-25* and *lin-45*), innate immune response (*atf-7, K09D9.1*, and *fbxa-150*), and hydrolase activity (*cnt-1, tbc-9, hgap-1*, and *Y92H12BM.1*). While genes in module 4 showed gradually decreasing expression levels with increasing regrowth rates, expression of genes in modules 5 and 6 showed a steep decrease at 2 and 6 µm/h, respectively. Interestingly, genes in these modules had minimal overlap with the DEGs identified when comparing regrowing and uninjured neurons in the previous section (Fig. 3i). GO analysis of all modules indicated a role of calcium signaling pathway, corroborating with prior findings^8^ (Supplementary Fig. 10). This result highlights the power of fs-LM to enable novel studies by facilitating isolation of cells on the basis of their phenotype.

## Discussion

In this work, we present fs-LM as a new method for isolating cells of interest on the basis of specific phenotype. The high precision of fs-laser ablation minimizes collateral and out-of-focus damage, allowing us to resect intact single cells directly from living tissue. As a demonstration, we isolated PLM neurons of 2 – 3 µm in diameter from living *C. elegans*. By isolating intact living neurons, we were able to achieve sensitive and specific gene expression profiling by scRNA-seq. More importantly, we show that fs-LM can minimize stress to the cells being isolated, thus avoid transcriptional artifacts while attaining a higher reproducibility in scRNA-seq compared to the dissociation-FACS method.

With these unique advantages, fs-LM can overcome many technical limitations inherit to the state-of-the-art cell isolation methods. The prevalent dissociation-FACS method induces stress related transcriptional artifacts in isolated cells. Its low cell yield and requirement for fluorescence tagging also limit its application to novel or rare cell populations. Most importantly, contextual and phenotypic information of the isolated cells are often lost during tissue dissociation. LCM can retain such information, but it requires a series of tissue processing steps (freezing, thin slicing, fixing, staining, dehydration. etc.) that pose high risk of degradation. Thin slicing of the tissue may also fragment cells of interest. Patch-seq is also prone to similar risks^36^, as aspirated cytosol can become degraded or contaminated inside the micropipette. As for purification-based isolation methods, such as transcriptome *in vivo* analysis^68^ and translating ribosome affinity purification^69^, only part of cellular content can be captured, which will likely suffer additional loss during column purification. Recent years have seen rise of *in situ* sequencing methods^70, 71^. fs-LM is complementary to such methods as they are still limited to mRNA, while intact cells isolated by fs-LM is amenable to all kinds of assays. In terms of limitations, fs-LM has a relatively low throughput compared to the dissociation-FACS method, which makes it more suitable for precision and phenotype-driven studies.

To demonstrate the capability of fs-LM to facilitate previously impossible studies, we isolated single regenerating PLM neurons from *C. elegans* for scRNA-seq, which has not been possible due to a lack of cell isolation methods. We uncovered a total of 1,217 DEGs whose selective inductions lead to successful or failed axon regrowth following laser axotomy, including 48 known regeneration regulators. Screen for upstream regulators of these DEGs identified 38 transcription factors as potential regeneration regulators. Heterogeneity of gene expression among seemingly homogeneous cell population has never been explored in the context of nerve regeneration. With fs-LM, we isolated single PLM neurons on the basis of axon regrowth rates and identified 226 genes with correlating expression levels. WGCNA revealed that the heterogeneity in axon regrowth rate was mediated by 6 gene modules with functions such as neurogenesis, RNA metabolism, DNA repair, and immune response. Our findings not only provide new insights into the known regeneration mechanisms in *C. elegans*, but also offer many new candidates for future studies.

With complete blueprint of anatomy, wiring diagram, and function, the *C. elegans* nervous system has long been one of the most well-defined models in neurology. Recently, its molecular topography has also been completed by the CeNGEN project^72, 73^. Combining such comprehensive knowledge base and fs-LM’s unique capabilities will finally allow mechanisms of development, regeneration, and degeneration to be studied in relation to neuronal genetic composition and physiology in a complete nervous system. Beyond *C. elegans*, fs-LM can also be readily applied to other living model organisms with reasonable modifications. As a demonstration, we applied fs-LM to isolate single cortical neurons from acute mice brain slices (Supplementary Fig. 11). In summary, our work establishes fs-LM as a versatile single cell isolation method with unique capabilities to facilitate future studies.

## Methods

### *C. elegans* strains and maintenance

We used the following strains in this work: CZ10175 *zdIs5 [Pmec-4::GFP + lin-15(+)]*, CZ11327 *dlk-1(ju476*); *zdIs5*, and N2. Both CZ10175 and CZ11327 strain express GFP in the 6 touch receptor neurons. *C elegans* were grown and maintained on nematode growth media (NGM) plates with bacteria of strain HB101 at 20 °C as previously described^5^. To obtain age-synchronized populations for FACS experiments, we lysed gravid adults with alkaline hypochlorite solution and collected the eggs in M9 buffer. The eggs were suspended in M9 buffer and allowed to hatch overnight on a rotor for aeration. The hatched larvae were arrested in the larval stage 1 (L1) until food was introduced. The synchronized L1 population was plated on NGM plate and allowed to grow for 50 – 52 hours until ready for harvest at middle to late larval stage 4.

### Laser axotomy

Femtosecond laser axotomies were performed on *C. elegans* anesthetized on agar pads. Agar pads were prepared by sandwiching 0.25 mL of melted 2% agar between two microscope slides. For each batch of axotomy, we used 15 µL of 10 mM levamisole solution to immobilize 8 – 12 L4 stage animals under a coverslip on the agar pad. Axotomy was performed on an upright microscope using a 60x, 1.4 NA oil immersion objective (Olympus LUMPFLFLN60XW). We ablated the axon of the PLM neuron that was closer to the coverslip at 50 µm from the soma. For ablation, we used a train of 300 laser pulses with 7.5 nJ pulse energies that were generated from an amplified femtosecond laser system (Spectra Physics “Spitfire”, 805 nm center wavelength, 250 fs pulse width, 1kHz repetition rate). We recovered the axotomized animals in a small droplet of M9 buffer and placed on the bacteria lawn of a fresh NGM plate. The axotomized animals were allowed to recover at 20 °C prior to imaging or fs-LM. From immobilizing the first animal of the batch to collecting the last axotomized animal from the agar pad, each batch of axotomy was consistently completed within 7 – 8 minutes to avoid stressing the animals. For axon regrowth length measurements, we took z-stack images (2 µm spaced) of the regenerating axons. Length measurement was taken on a max-projection image formed from the z-stack images by ImageJ. Only processes regrowing longer than 5 µm were included in length measurements. We excluded reconnected axons from the statistical analysis. For mock axotomies, ablation was performed on the hypodermis instead of the axon (at 2 µm from the position where actual axotomies would be performed).

### *C. elegans* dissociation and flow cell activated sorting (FACS)

For dissociation-FACS isolation of *C. elegans* touch receptor neurons, we followed an established protocol for dissociation of *C. elegans^32^*. We collected a synchronized animal population with M9 buffer from NGM plates and washed the animal pellet with egg buffer 3x to remove bacteria. The pellet was then incubated in a freshly prepared 0.25% sodium dodecyl sulfate, 200 mM dithiothreitol solution for 4 minutes to weaken the cuticle of the animals. We washed the treated pellet 5x with egg buffer, then suspended the pellet in freshly prepared 15 mg/mL Pronase solution to further digest the cuticle for 20 minutes. We gently pipetted the entire suspension up and down with syringe and 18G needle to promote digestion and release of cells. During pipetting, bubbles were carefully avoided. At the end of the Pronase treatment, we stopped reaction by adding pre-chilled L15 medium supplemented with 10% feral bovine serum (FBS) and allowed the undigested carcass to settle at 4 °C for 5 minutes. The cell suspension remained chilled from this point forward. We collected the supernatant and washed the collected cells 2x with L15 containing 2% FBS. The final concentration-adjusted suspension was filtered through a 5 µm syringe filter. Prior to FACS, we added propidium iodide (PI) to the cell suspension for live/dead staining. FACS was performed using a BD FACSAria II sorter. The GFP sorting gate was determined using cell suspension collected from N2 animals as reference, and the PI sorting gate was determined by test sorting a stained cell population and observing separation of live/dead cells into two clusters. The GFP+ PI-events were sorted directly into a 96-well plate (single neuron sample, n = 26) or 1.5 mL tubes (bulk sample each containing > 5,000 single neurons, n = 3) containing lysis buffer with 2 U/µL RNAse inhibitor. The collected GFP+ PI-neurons were 0.3% of all sorted cells. We confirmed purity of the collected neurons by sorting extra GFP+ PI-events into L15 medium and visually inspecting the collected cells under a microscope. The sorted samples were promptly frozen on dry ice for subsequent reverse transcription of cDNA.

### fs-LM of *C. elegans* PLM neurons

We started by taping a clean 25 x 75 mm coverslip to a microscope slide of the same size. The microscope slide provides mechanical support when mounted on microscope stage. Then we pipetted 4 droplets of buffers onto the coverslip: 25 µL of dissection buffer (1 mg/mL Pronase in egg buffer), 25 µL of L15 medium with 30% FBS, and 2 x 25 µL fresh L15 medium. Prior to fs-LM, we washed each batch of 4 animals in M9 buffer to remove bacteria and anesthetized them using 20 mM levamisole for 4 minutes until movements were suppressed. The anesthetized animals were then transferred into the dissection buffer using an eyelash pick. We mounted the slide onto an upright microscope, equipped with XYZ motorized stage (ASI PZ-2000FT). fs-LM was subsequently performed using a 2 mm working distance, 60x, 1.0 NA water dipping objective (Olympus LUMPLFLN60XW). We first resected the target neuron by ablating its surrounding tissue at a series of spots arranged in a hemispherical pattern encasing the neuron (as discussed in the main text). Each ablation spot was generated by a train of 300 laser pulses with 16 nJ pulse energy, generated from the same laser as used for laser axotomy. We slightly increased pulse energy to 20 nJ when ablating spots deeper underneath the cuticle to compensate for scattering. Towards the end of the resection process, axon of the target neuron was ablated at a distance of 20 µm from the neuronal soma. For puncturing the cuticle, 150 laser pulses with 50 nJ pulse energy were used. The resected neurons would normally be released through the incision driven by internal pressure of the animal, which could also be assisted by repeatedly delivering 150 laser pulses with 50 nJ pulse energy to a spot ~50 µm away from the neuron, thus generating a cavitation bubble that can push the neuron towards the incision. In case the released neuron was stuck to debris, it could be separated by inducing laser bubbles in the buffer at ~50 µm from the neuron, which could generate torrent in the buffer that effectively breaks up the debris without damaging the neuron.

To aspirate the released neuron, we used a glass micropipette controlled pneumatically by a microinjector (Narishige IM-11-2), which was mounted on a 3-axis micromanipulator (Narishige MN-153). The micropipettes had a tip inner diameter of 10 – 15 µm and were polished by a microforge. We coated the inner surface of the micropipette with silicone (Sigmacote, Sigma-Aldrich). Prior to fs-LM, we filled the tip of the micropipette with 30% FBS/L15 buffer, positioned the tip at the center of field of view, then retracted the tip so that during fs-LM the tip would not be in contact with the dissection buffer droplet. When a neuron was successfully released, we extended the tip while applying low positive pressure to generate a slight outflow. We used the outflow to wash and elevate the released neuron away from other cells or debris, after which we aspirated and collected the neuron from the dissection buffer droplet. The collected neuron was next washed 1x in the 30% FBS/L15 buffer droplet and 2x in fresh L15 medium droplets. For each wash, we completely flushed the micropipette (without creating bubbles) to guarantee cleanliness. After the last wash, we pipetted 6 µL of 1x lysis buffer prepared from Clontech SMART-seq v4 kit with 2 U/µL RNAse inhibitor onto the coverslip. The collected neuron was then deposited into the droplet of lysis buffer, which we aspirated and transferred to a PCR tube for freezing at −80 °C. Identical fs-LM protocol was applied to collect uninjured wild-type (n = 23), uninjured *dlk-1 (0)* (n = 25), axotomized wild-type (n = 47), and axotomized *dlk-1 (0)* (n = 25) PLM neurons. For each animal, only one neuron was isolated by fs-LM.

### Single cell RNA-sequencing and data analysis

cDNA libraries were constructed with Clontech SMART-seq v4 3’ DE Kit per manufacturer’s instruction. cDNA libraries were pooled and NGS libraries were constructed by Illumina Nextera XT DNA Library Prep Kit. We performed pair-end 150 bp sequencing with Illumina Hi-seq 4000. The raw sequencing data was demultiplexed and filtered to remove low-quality reads by Cutadapt v2.8. We aligned the filtered reads to *C. elegans* reference genome WS270 using STAR aligner v2.7.2. The alignments were summarized using the featureCount function from Subreads package v1.6.4. Only unique alignments were retained. Each individual neuron received over 3 million summarized counts, which was deep enough to maximize gene discovery according to subsampling analysis. As Clontech SMART-seq v4 3’ DE Kit sequences the 3’ end of amplified cDNA fragments, we used CPM as normalized metric of gene expression. For differential expression analysis, we used single cell differential expression package v3.12. For detailed description of data analysis discussed in the paper, please refer to Supplementary Methods.

## Contributions

A.B. conceived the method, and supervised the overall direction of the research. P.Z., C.M., and A.B. designed the experiments. P.Z. and C.M. further developed the method to bring it into practice, the optical setup, and the automated LabView program. P.Z. maintained animal strains, performed laser axotomy experiments, and isolated neurons used in this study. P.Z. performed RNA-sequencing and data analysis with inputs from K. Ma. P.Z. and A.B. prepared the manuscript with inputs from all authors.

## Data availability

The ChIP-seq datasets used for identifying potential regulators of nerve regeneration are publicly available at https://www.encodeproject.org/ (ENCODE Project^74, 75^). All RNA-sequencing data involved in Fig. 1–3 are available on NCBI SRA repository under study identifier SRP300789.

## Acknowledgements

This work was supported by the National Institutes of Health (NIH) Grant no. R21-NS109821 and by a grant from The University of Texas System Neuroscience and Neurotechnology Research Institute (UTS-NNRI). We are grateful to R. Maiya and R. Messing (The University of Texas at Austin) for sharing insights and brain slice samples and equipment during the early-stage development of this method. We acknowledge Y. Li (Dell Pediatric Research Institute) for assistance during FACS experiments, Y. Jin (University of California San Diego) for *C. elegans* strains used in this study (CZ10175 and CZ11327), W. Chad (The University of Texas at Austin) for preparing micropipette, and S. Mondal and E. Hegarty (The University of Texas at Austin) for help with animal maintenance and insights in experimental design. We thank the ENCODE Consortium, in particular M. Snyder, V. Reinke and K. White for providing the ChIP-seq datasets. We thank the Caenorhabditis Genetic Center (CGC) for providing us with N2 animals for the study.

---

**Supplementary Fig. 1.**
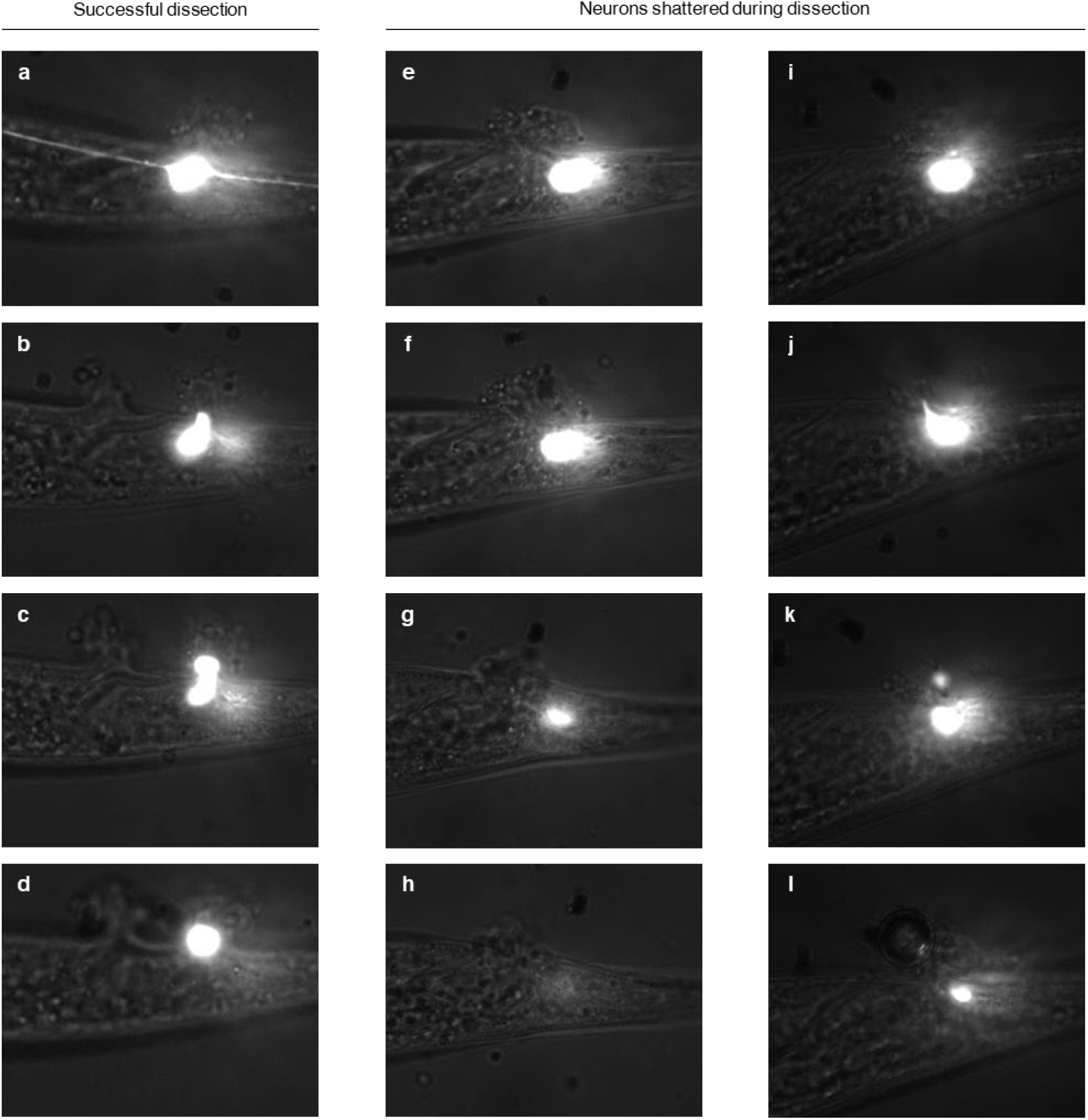
Successful and unsuccessful fs-LM isolation of *C. elegans* PLM neurons. **a-d,** Example of successful fs-LM isolation of neurons. The resected neuron was released through the incision smoothly without damage, as evidenced by consistent volume and cytoplasmic GFP intensity. **e-h**, Incomplete resection of neuron causing it to remain in place after the incision was made. As a result, the neuron was torn by outflow of surrounding tissue. **i-l,** A resected neuron was fragmented when migrating through a small cuticle incision. In this case, the neuron was moving towards the incision, which indicated complete resection.

**Supplementary Fig. 2.**
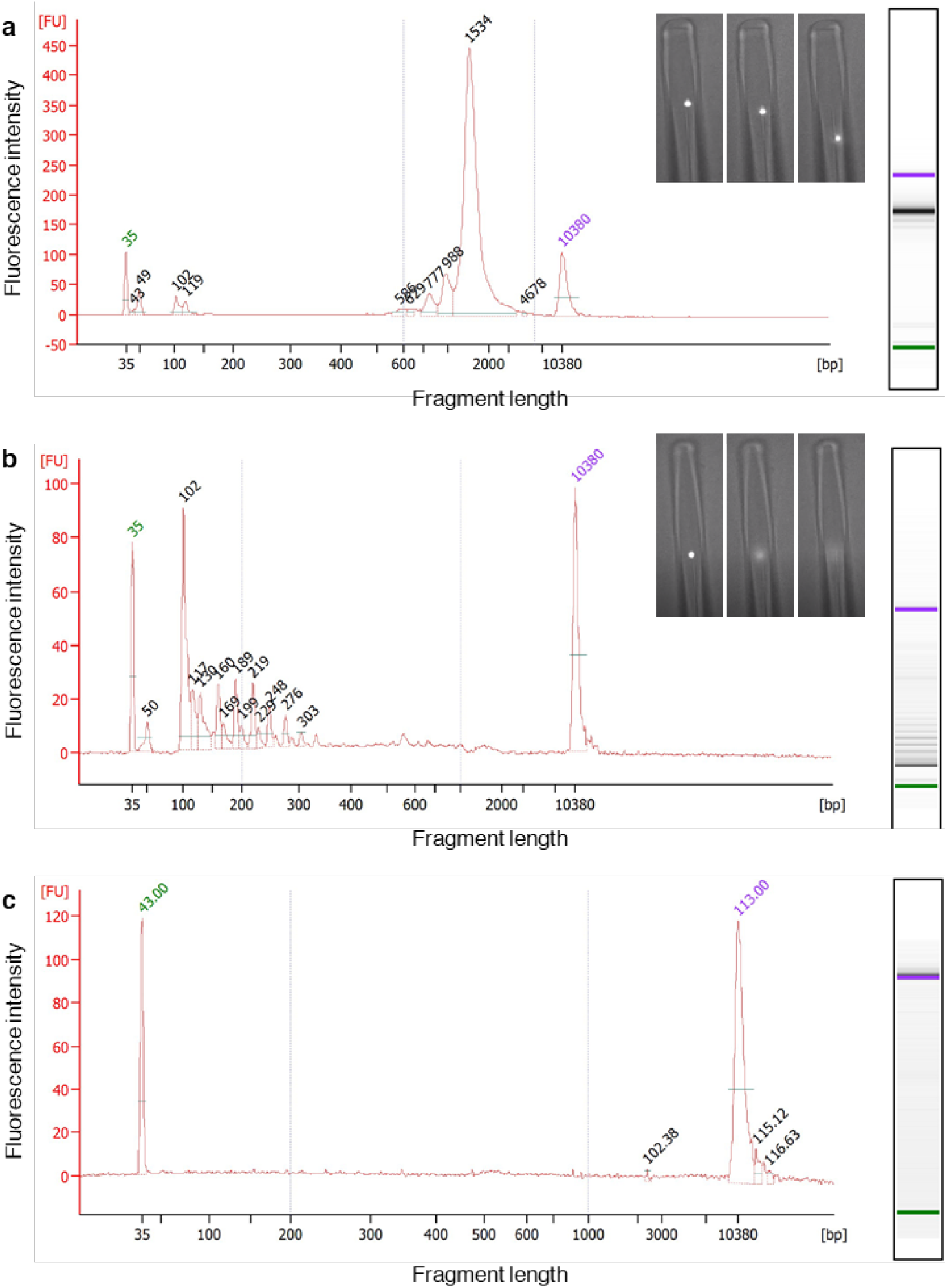
Quantification of amplified cDNA libraries from fs-LM isolated single *C. elegans* neurons. **a,** Example of libraries with good quality. Inset: the isolated neurons retained GFP intensity until being deposited into lysis buffer. **b**, Example of libraries that failed quality check, and were thus excluded from this study. Inset: the isolated neuron lysed before being deposited into lysis buffer. Although the cellular content of the neuron supposedly remained in the micropipette, degradation and attachment to inner wall of the micropipette severely affected RT-PCR, resulting in libraries dominated by short fragments that were mostly primer-dimer. **c,** Example of NTC controls. No amplified cDNA were detected as expected.

**Supplementary Fig. 3.**
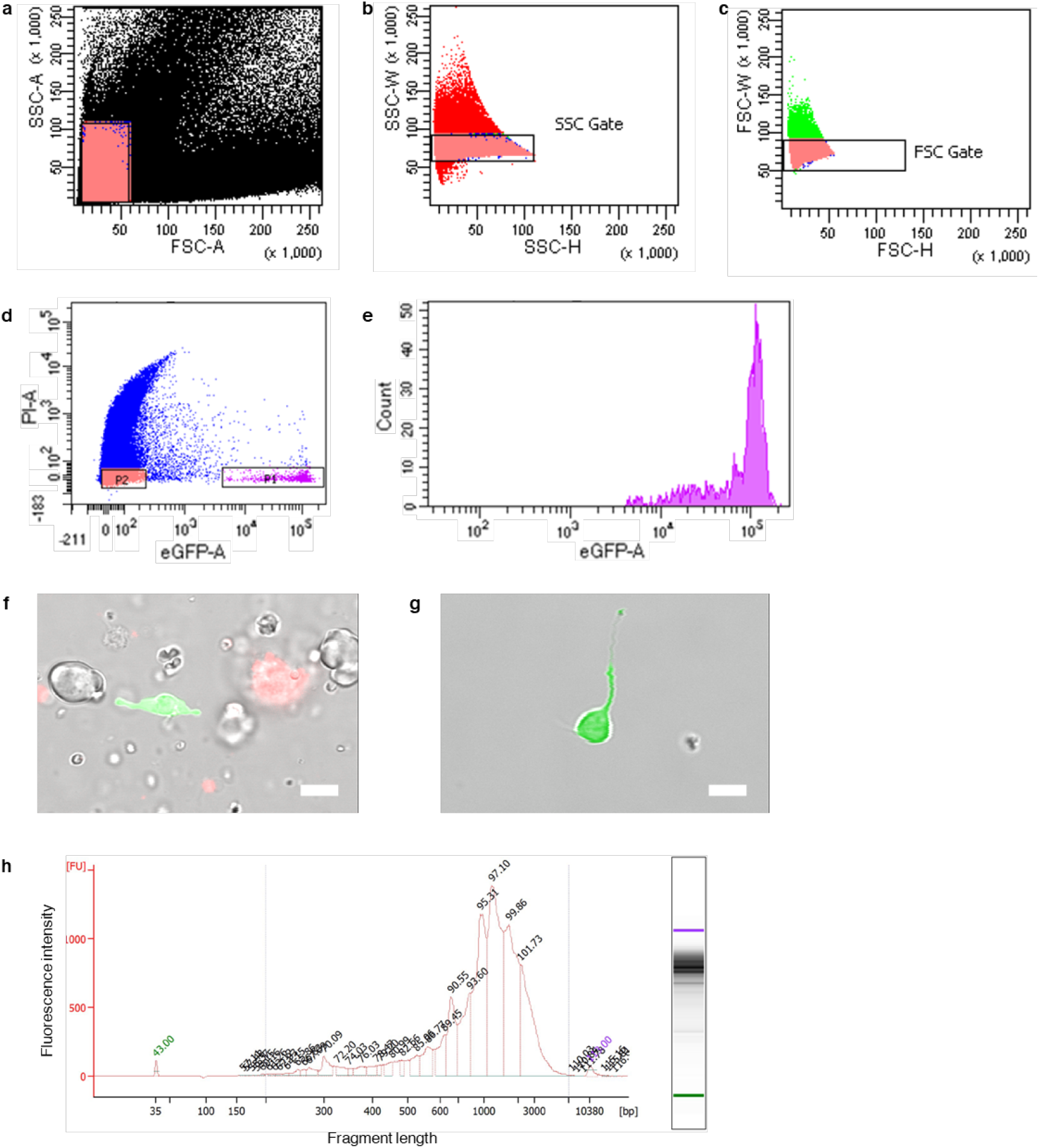
Isolation of *C. elegans* touch receptor neurons by the dissociation-FACS method. **a-d,** FACSAria II gate settings for isolating living GFP-labeled *C. elegans* touch receptor neurons from dissociated animals. Prior to FACS, the cell suspension was filtered through a 5 µm syringe filter to prevent clogging. SSC-A, SSC-W, and FSC gates were set up to exclude debris and clusters of cells. We stained the dead neurons with propidium iodide. GFP+ PI-events were sorted into 96 well plates or 1.5 mL conical tubes containing lysis buffer. The GFP threshold was determined in reference to the autofluorescence of cells from dissociated N2 worms. The PI threshold was determined by test sorting a small number of PI-stained cells and observing two distinct population of live/dead cells. **e,** Histogram of GFP levels of the isolated neurons. **f**, Cell suspension prior to sorting and **g,** example of collected neuron. Green: GFP. Red: PI. Scale bar: 5 µm.

**Supplementary Fig. 4.**
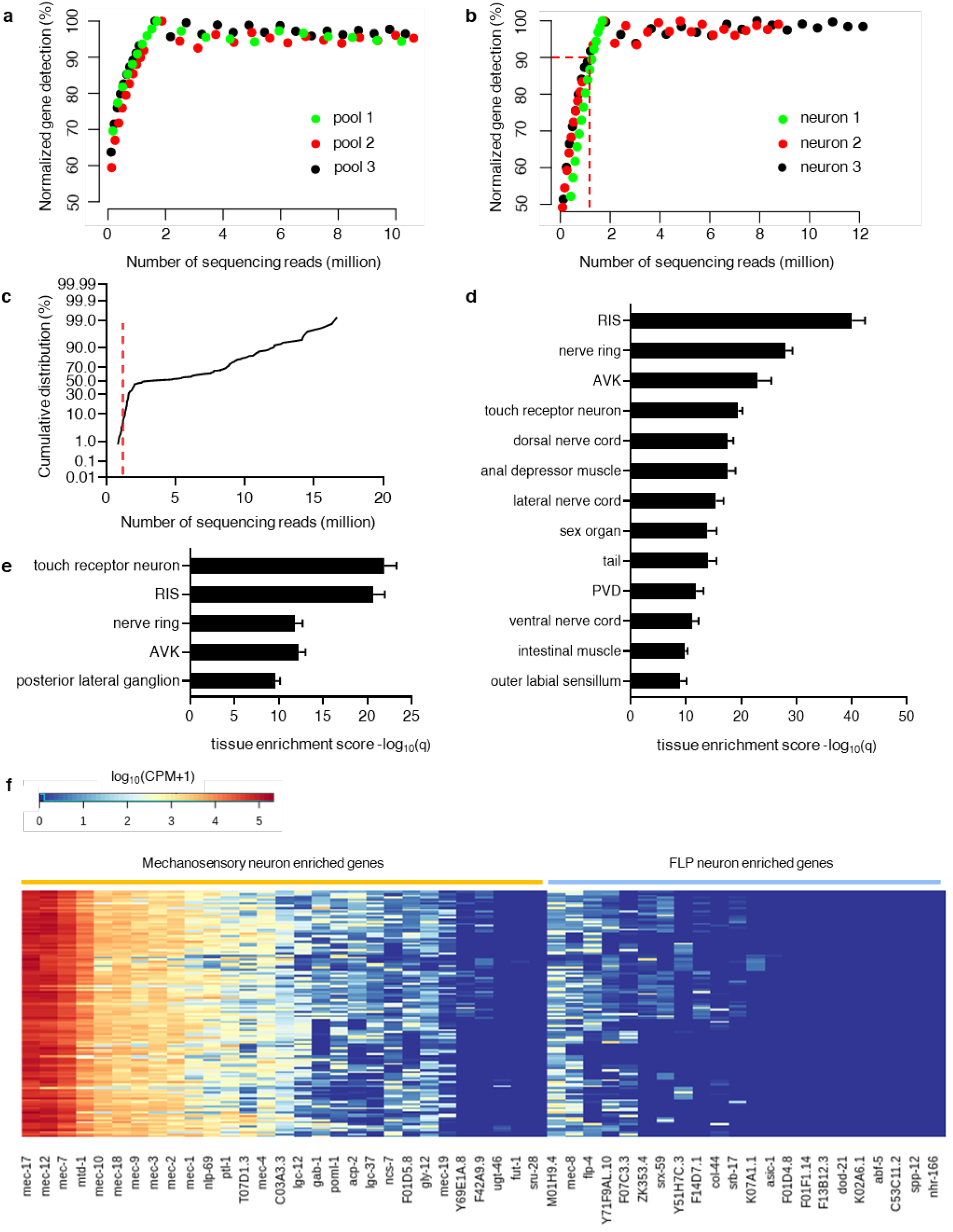
RNA-sequencing of fs-LM and dissociation-FACS isolated neurons. **a, b,** Sufficient sequencing depth was confirmed by subsampling reads from pooled and single neurons samples and checking the numbers of genes detected. We found that by sequencing neurons at 2 million uniquely mapped reads per sample, the numbers of genes detected reached plateau. We discarded all single neuron samples with less than 1 million uniquely mapped reads to avoid technical bias induced by RNA-sequencing. **c,** Cumulative distribution of sequencing depth among all samples. Less than 10% of single neuron samples were discarded due to insufficient sequencing depth. **d, e,** Top tissue enrichment analysis^18^ terms of dissociation-FACS isolated single neurons (d) and fs-LM isolated single neurons (e). fs-LM isolated neurons correctly showed touch receptor neuron as the top term, while dissociation-FACS isolated neurons showed RIS, nerve ring and AVK as the top 3 terms, likely due to collection of all 6 touch receptor neurons. **f,** Expression levels of marker genes among fs-LM isolated single neurons. In the *C. elegans* strain we used (CZ10175 and CZ11327), all 6 touch receptor neurons were labeled by GFP. These touch receptor neurons include 4 mechanosensory neurons (PLML, PLMR, ALML, ALMR) and 2 FLP interneurons (AVM, PVM). Genes specifically upregulated in mechanosensory and FLP neurons have been previously characterized^19^. Because fs-LM specifically isolated PLM neurons, higher expression levels were found among genes enriched in mechanosensory neurons, while expression levels of genes enriched in FLP neurons were low.

**Supplementary Fig. 5.**
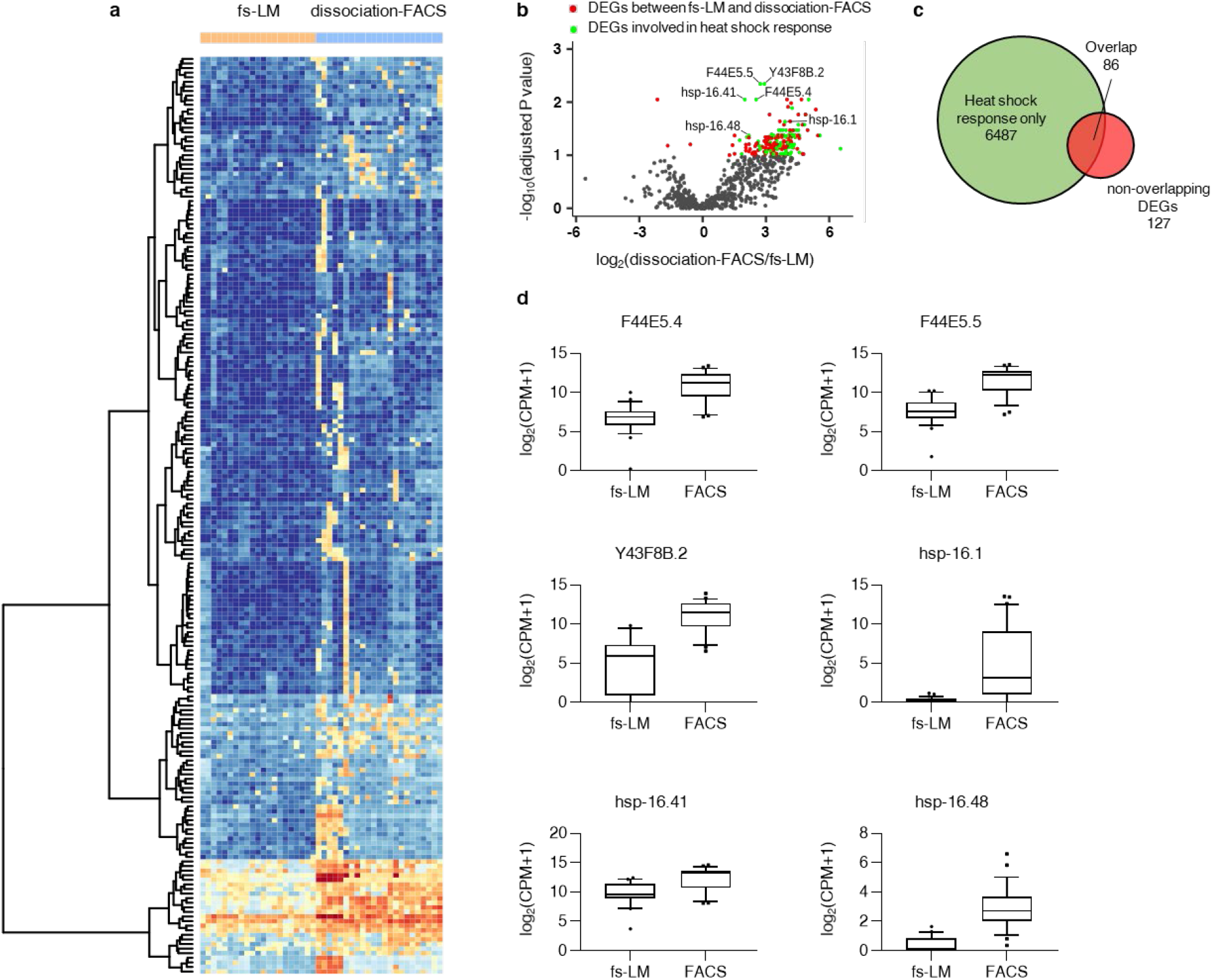
Differential gene expression between neurons isolated by fs-LM and dissociation-FACS method. **a,** Heatmap showing expression levels of the DEGs (FDR < 0.1) in neurons isolated by fs-LM and dissociation-FACS method. **b,** Up-regulated genes in dissociation-FACS isolated neurons showed significant overlap with genes up-regulated during heat shock response. **c,** Overlap between DEGs and heat shock response genes^20^, indicating a significant correlation between heat shock response and gene regulatory pattern displayed by the dissociation-FACS isolated neurons (Chi-square test, *P* = 0.023). **d,** Gene expression levels of highly up-regulated DEGs, which were also highly upregulated during heat shock response dependent on heat shock factor 1.

**Supplementary Fig. 6.**
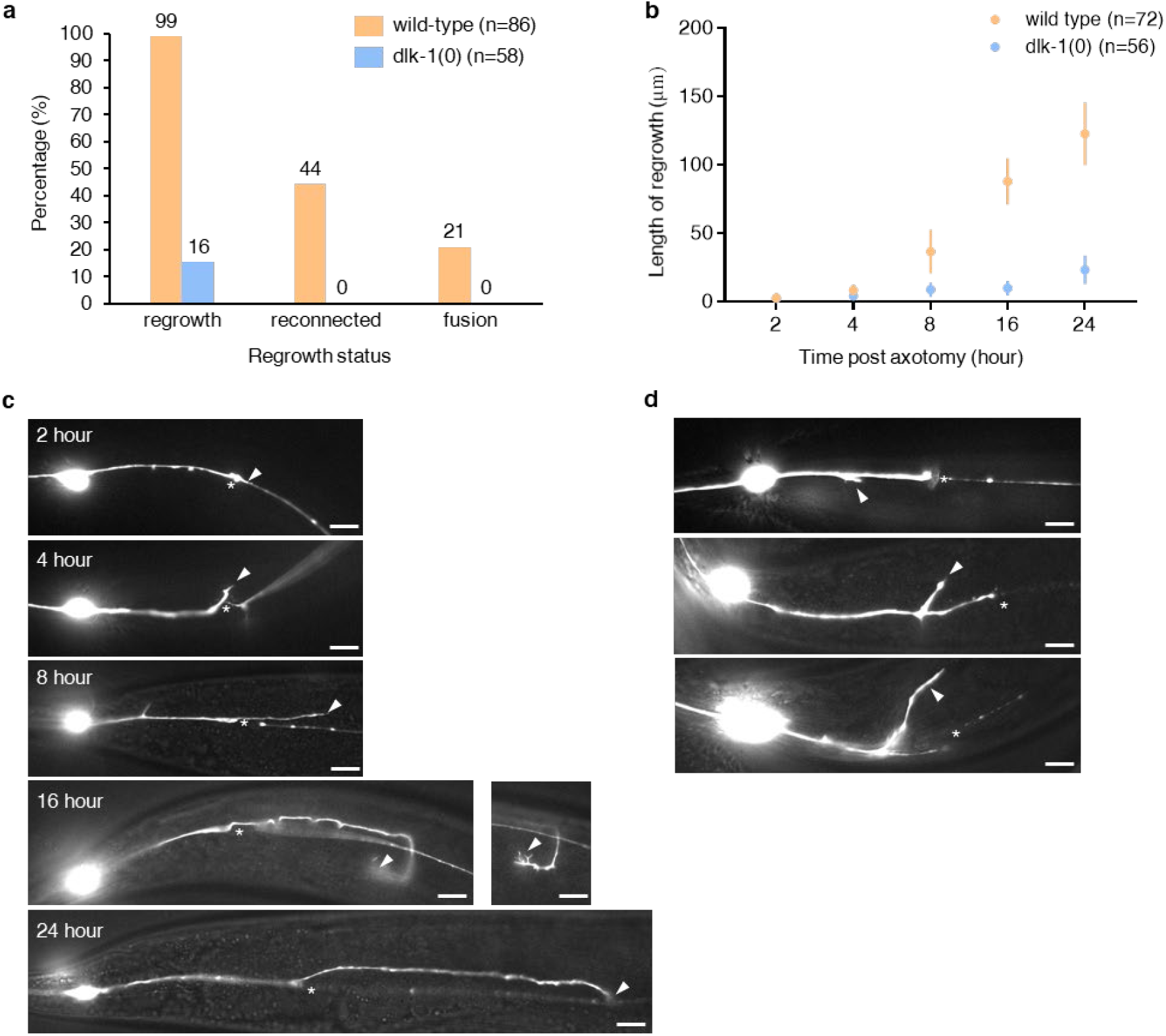
Axon regeneration of PLM neurons in wild-type and *dlk-1 (0)* animals following fs-laser axotomy. **a,** Regeneration status of axotomized PLM neurons at 24-hour post axotomy. Only 1 out of 86 wild-type PLM neurons failed to initiate regrowth, likely due to improper laser axotomy inducing excessive damage. Among the PLM neurons that regrew, 44% and 21% reconnected and fused respectively with the distal end, which was consistent with our previous findings^5^. Interestingly, we also observed ectopic regeneration in a small number of *dlk-1 (0)* animals, which can be attributed to DLK-1 independent regeneration pathways^7^ and incomplete *dlk-1* suppression. **b,** Length of axon regrowth at various time points post axotomy. At 2-4 hour post axotomy, wild-type PLM neurons started to regrow at an average rate of 6.1 µm/hour. The ectopic regrowth rate of *dlk-1 (0)* neurons was found to be 1.2 µm/hour. **c, d,** Microscope images of regrowing wild-type (c) and *dlk-1 (0)* neurons (d). Wild-type neurons initiated regrowth from the proximal stump led by growth cone, while *dlk-1 (0)* neurons regrew mostly by sprouting from the axon instead of the proximal stump. No obvious growth cone was observed. Asterisk, site of laser axotomy. Arrowhead, tip of regrowing axon. Scale bar, 10 µm.

**Supplementary Fig. 7.**
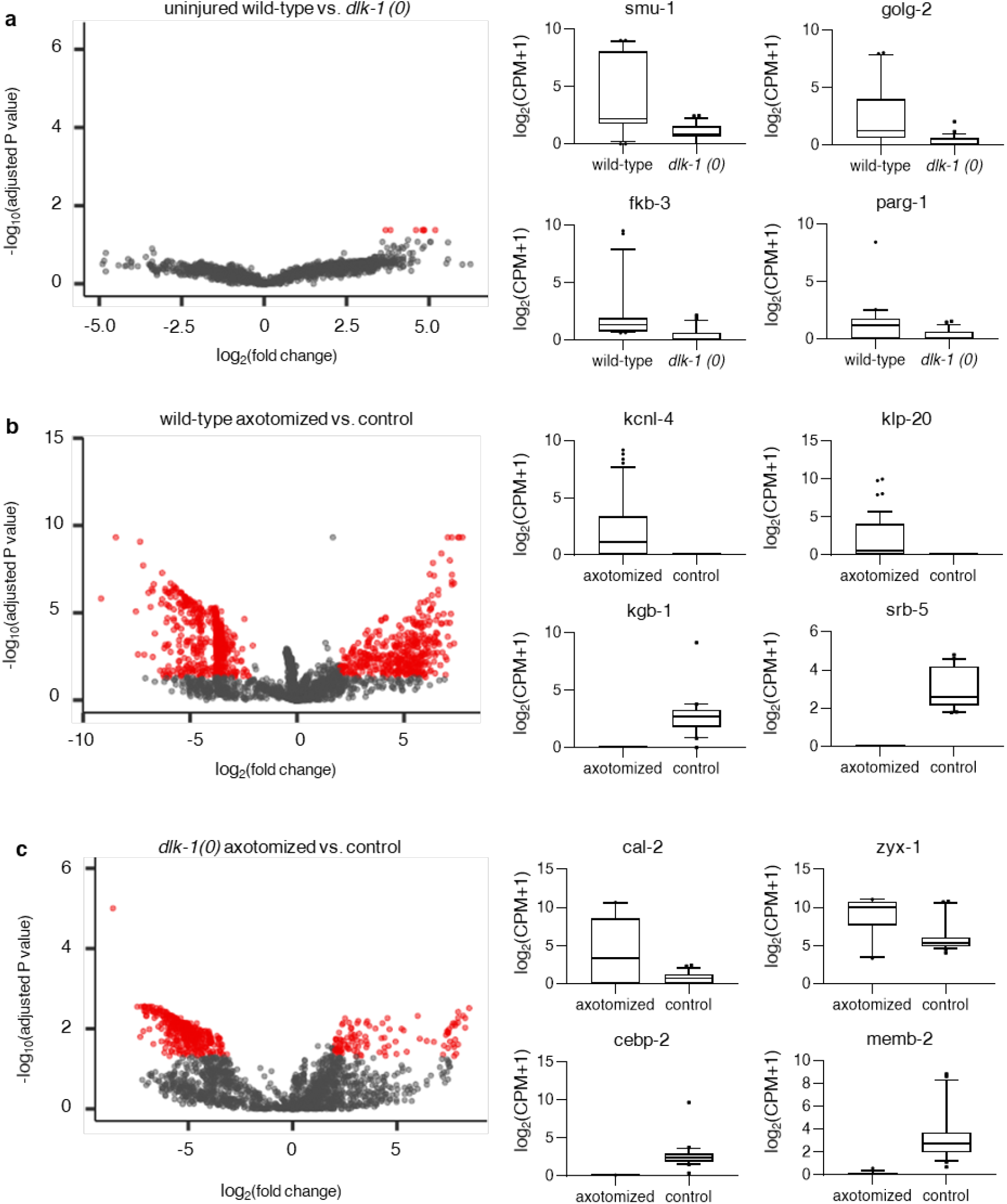
Differential gene expression between uninjured/axotomized neurons isolated from wild-*type/dlk-1 (0)* animals. **a,** DEGs between uninjured wild-type and *dlk-1 (0)* neurons. Red points in volcano plot represent significant DEGs with adjusted *P* value < 0.05 and fold change > 2. Boxplots show expression levels of top DEGs. *smu-1* encodes an ortholog of vertebrate SMU protein. *golg-2* encodes an ortholog of human GOLGA2. *fkb-3* encodes a peptidylprolyl cis/trans isomerase regulated by DAF-2 insulin receptor-like pathway and DAF-16 transcription factor. *parg-1* displayed a higher average expression level in wild-type neurons, which is consistent with prior finding^21^, although the change was not significant. **b,** DEGs between uninjured and axotomized wild-type neurons. *kcnl-4* encodes an ortholog of human KCNN1 and KCNN4 (potassium calcium-activated channel subfamily N member 1/4). *klp-20* encodes a kinesin-like protein and exhibits microtubule motor activity. *kgb-1* is a regulator of nerve regeneration and encodes a member of the JNK subfamily of MAP kinases. *srb-5* encodes a predicted serpentine receptor. **c,** DEGs between uninjured and axotomized *dlk-1 (0)* neurons. *cal-2* encodes a calmodulin homolog and is predicted to have calcium ion binding and enzyme regulator activity. *zyx-1* encodes a LIM domain protein similar to vertebrate Zyxin with predicted role in anchoring actin filaments to dense bodies. *cebp-2* encodes a predicted transcription factor and is involved in defense response to bacteria. *memb-2* encodes an ortholog of human GOSR2 and is predicted to have SNAP receptor activity and SNARE binding activity.

**Supplementary Fig. 8.**
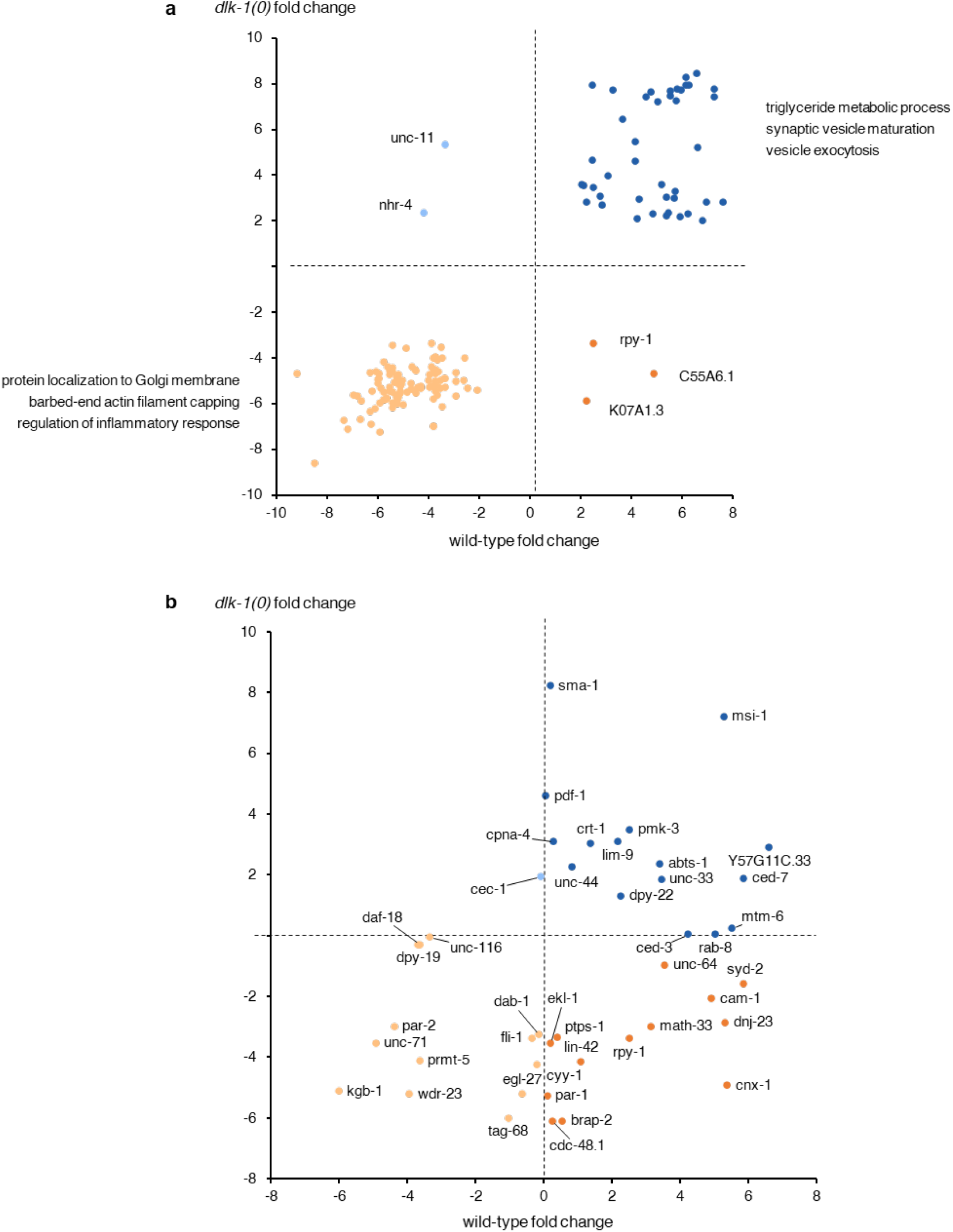
Pattern of differential expression displayed by specific groups of DEGs. **a,** Differential expression pattern of DEGs between axotomized and uninjured neurons shared by wild-type and *dlk-1 (0)* animals. Most of these shared DEGs displayed concordant (Pearson’s correlation coefficient = 0.94, *P* = 2 x 10^−16^) differential expression patterns in neurons from wild-type and *dlk-1 (0)* animals. Gene Ontology (GO) analysis of up-regulated and down-regulated DEGs identified functions such as vesicle maturation, exocytosis, and inflammatory response, which likely reflects the neurons’ response to injury rather than induction of axon regeneration. **b,** Differential expression pattern of known regeneration regulators. These regulators exhibited a variety of differential expression patterns, except for down-regulated in wild-type neurons and up-regulated in *dlk-1 (0)* neurons. This is likely because DEGs in *dlk-1 (0)* animals were mostly down-regulated following axotomy. Note that change in expression levels is not necessarily part of the known regulatory mechanisms of these regulators. For example, KGB-1 regulates regeneration through kinase activity as part of DLK-1 signaling pathway. The detected differential expression patterns of these regulators can provide new insights into their regulatory mechanisms or hint potential co-factors.

**Supplementary Fig. 9.**
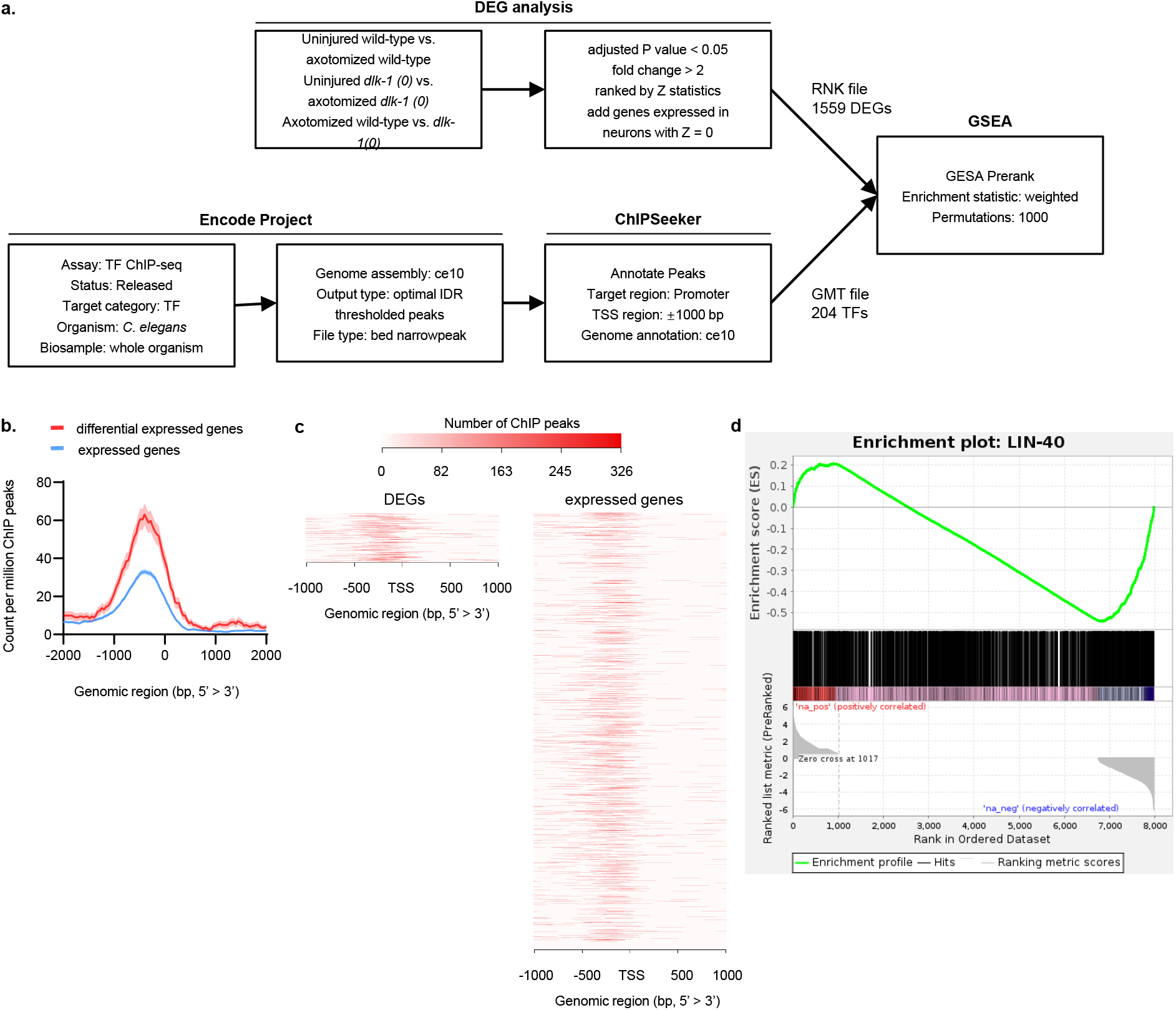
Screening for upstream regulators of the identified DEGs based on transcriptional factor binding affinity. **a,** Data sourcing, processing, and analysis to identify *C. elegans* transcription factors (TFs) with potential regulatory roles in axon regeneration. Through these steps, we aimed to identify TFs with significantly higher binding affinity to promoter regions of the identified DEGs using all genes expressed in neuron as a benchmark. Statistical analysis was performed by Gene Set Enrichment Analysis (GSEA) Pre-ranked module^22^. As the first input to GSEA Pre-ranked module, we generated a list of all genes expressed in neurons ranked by their Z statistics (generated during differential expression analysis, assigned a Z statistics of 0 if not a DEG). Since larger Z statistics (either positive or negative) indicated deeper involvement in axon regeneration, such ranking placed genes involved in axon regeneration to the two extremes of the list. As the second input to GSEA Pre-ranked module, for each TF, we identified genes whose promoter region contained ChIP-seq peaks using ChIPSeeker^23^ (TF target genes). Based on both gene lists, GSEA Pre-ranked module can compute an enrichment score (ES), which corresponded to a weighted Kolmogorov-Smirnov-like statistic. If overlaps between the TF target genes and the ranked gene list were located towards the extremes of the ranked gene list rather than evenly distributed, GSEA Pre-ranked module will produce a larger ES (either positive or negative). Finally, a *P*-value was calculated based on null distribution obtained by permutating the ranked gene list 5000x. **b,** Metagene binding profiles of LIN-40 centered at the transcription start site. Red traces represent density of ChIP peaks mapped to the DEGs. Blue traces represent density of ChIP peaks mapped to all genes expressed in neurons. **c,** Heatmap of LIN-40 ChIP-seq peaks centered at the transcription start site. **d,** Result of GSEA Pre-ranked analysis for LIN-40 (*P* < 0.001).

**Supplementary Fig. 10.**
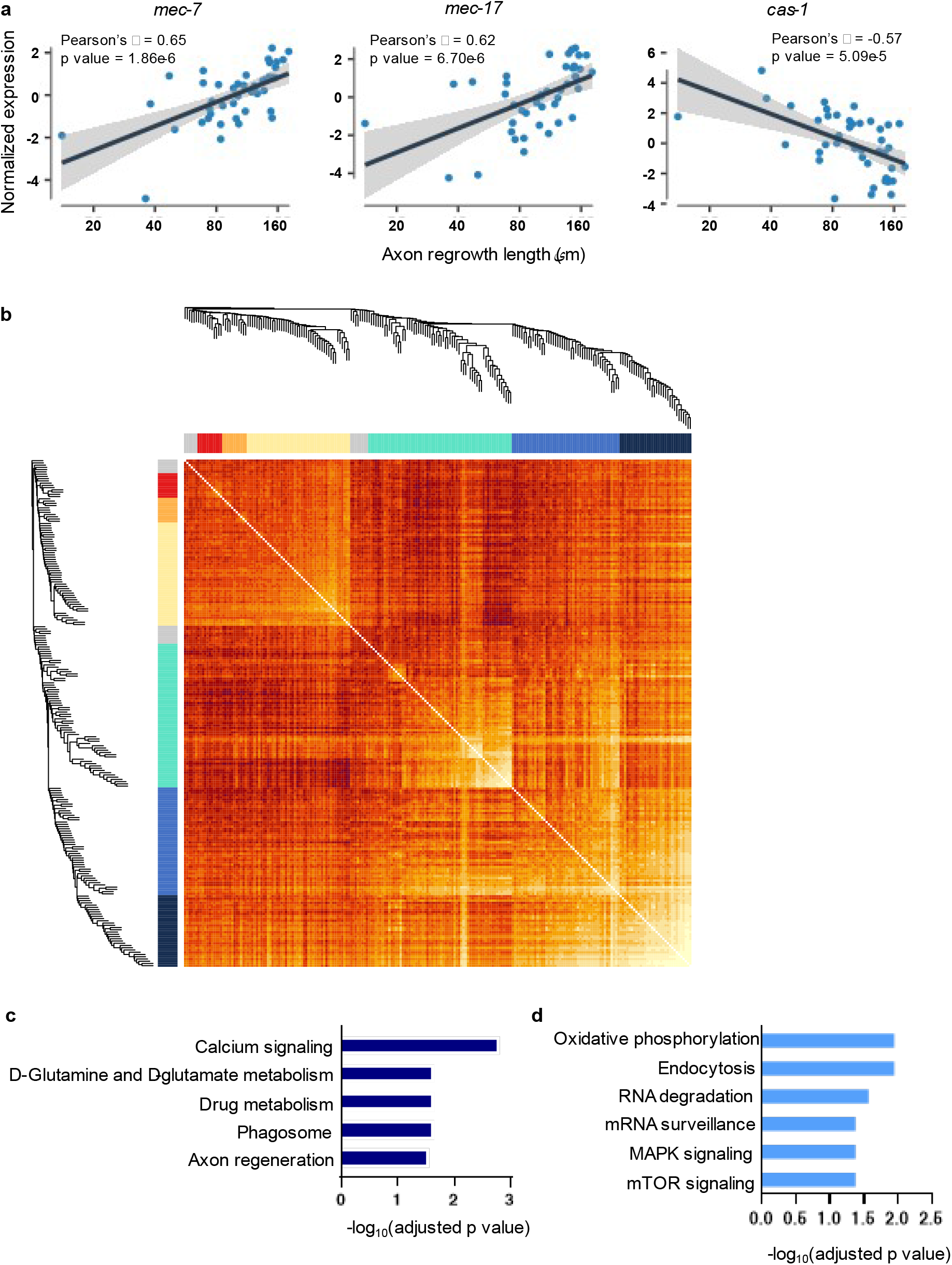
Weight gene co-expression network analysis of genes whose expression pattern showed correlation with axon regrowth rate across single regrowing neurons. **a,** Example of genes whose expression pattern showed significant correlation with axon regrowth rate across single regrowing neurons. Each plot shows expression pattern of the gene and results of linear fitting. Grey area represents 95% confidence interval. **b**, Inter-module correlation among the 6 gene modules identified by WGCNA. **c**, GO analysis for enriched KEGG pathways among all gene modules. **d**, GO analysis for enriched WikiPathways among all gene modules

**Supplementary Fig. 11.**
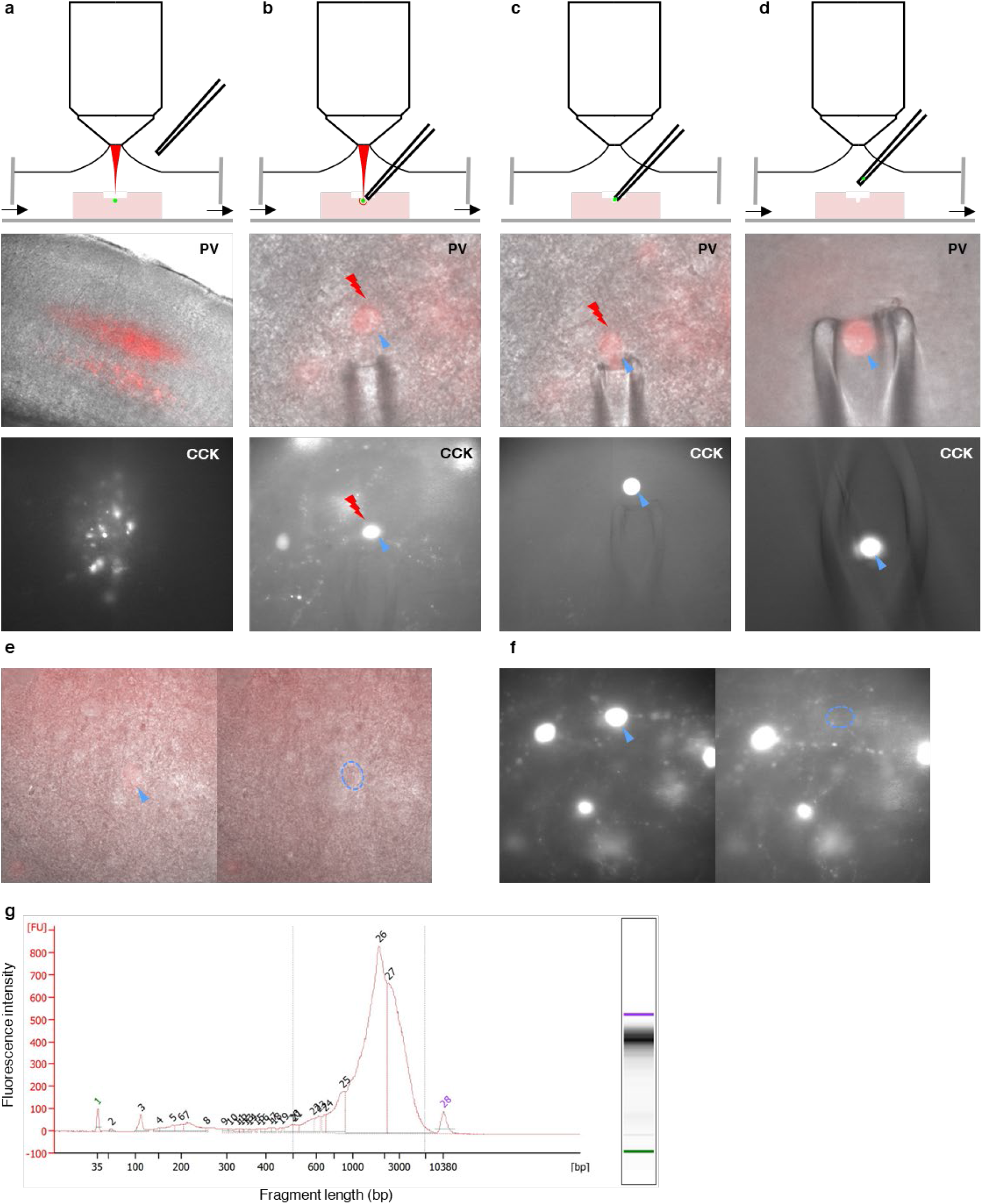
fs-LM isolation of single mice/gerbil brain neuron from acute brain slice. **a,** The 500 ptm thick acute brain slice was placed in a chamber and perfused with ice-cold, carbogen-infused artificial cerebrospinal fluid (aCSF). Prior to fs-LM, we removed the top layer of dead tissue using laser ablation to expose the target neuron. Microscope images show the brain region containing the target neurons (cortical PV neurons labeled by tdTomato and CCK neurons labeled by GFP in medulla). **b,** The micropipette was positioned close to the target neuron with a slight positive pressure applied to prevent contamination. We performed fs-LM as in *C. elegans* to resect the target neuron from surrounding tissue. **c,** Towards the end of fs-LM, the aCSF perfusion was paused in order to prevent the resected neuron from being carried away by liquid flow. Meanwhile, aspiration was applied from the micropipette. We stopped fs-LM as soon as the neuron started migrating towards the micropipette. **d,** The isolated neuron was cleaned in aCSF before being deposited into lysis buffer and frozen on dry ice.

## Notes

### Competing Interest Statement

The authors have declared no competing interest.

## References

1. Courtine, G. & Sofroniew, M.V. Spinal cord repair: advances in biology and technology. Nature Medicine 25, 898–908 (2019).

2. Hutson, T.H. & Di Giovanni, S. The translational landscape in spinal cord injury: focus on neuroplasticity and regeneration. Nature Reviews Neurology 15, 732–745 (2019).

3. Duan, X. et al. Subtype-Specific Regeneration of Retinal Ganglion Cells following Axotomy: Effects of Osteopontin and mTOR Signaling. Neuron 85, 1244–1256 (2015).

4. Norsworthy, M.W. et al. Sox11 Expression Promotes Regeneration of Some Retinal Ganglion Cell Types but Kills Others. Neuron 94, 1112–1120.e1114 (2017).

5. Yanik, M.F. et al. Functional regeneration after laser axotomy. Nature 432, 822–822 (2004).

6. Bourgeois, F. & Ben-Yakar, A. Femtosecond laser nanoaxotomy properties and their effect on axonal recovery in C. elegans. Opt Express 16, 5963–5963 (2008).

7. Hammarlund, M., Nix, P., Hauth, L., Jorgensen, E.M. & Bastiani, M. Axon Regeneration Requires a Conserved MAP Kinase Pathway. Scence 323, 802 (2009).

8. Ghosh-Roy, A., Wu, Z., Goncharov, A., Jin, Y. & Chisholm, A.D. Calcium and Cyclic AMP Promote Axonal Regeneration in Caenorhabditis elegans and Require DLK-1 Kinase. The Journal of Neuroscience 30, 3175 (2010).

9. Nix, P., Hisamoto, N., Matsumoto, K. & Bastiani, M. Axon regeneration requires coordinate activation of p38 and JNK MAPK pathways. Proceedings of the National Academy of Sciences 108, 10738 (2011).

10. El Bejjani, R. & Hammarlund, M. Notch Signaling Inhibits Axon Regeneration. Neuron 73, 268–278 (2012).

11. Zou, Y. et al. Developmental decline in neuronal regeneration by the progressive change of two intrinsic timers. Science 340, 372–376 (2013).

12. Byrne, Alexandra B. et al. Insulin/IGF1 Signaling Inhibits Age-Dependent Axon Regeneration. Neuron 81, 561–573 (2014).

13. Chuang, M. et al. The microtubule minus-end-binding protein patronin/PTRN-1 is required for axon regeneration in C. elegans. Cell Rep 9, 874–883 (2014).

14. Chen, L. et al. Axon injury triggers EFA-6 mediated destabilization of axonal microtubules via TACC and doublecortin like kinase. Elife 4, e08695 (2015).

15. Li, C., Hisamoto, N. & Matsumoto, K. Axon Regeneration Is Regulated by Ets–C/EBP Transcription Complexes Generated by Activation of the cAMP/Ca2+ Signaling Pathways. PLOS Genetics 11, e1005603 (2015).

16. Neumann, B. et al. EFF-1-mediated regenerative axonal fusion requires components of the apoptotic pathway. Nature 517, 219–222 (2015).

17. Alam, T. et al. Axotomy-induced HIF-serotonin signalling axis promotes axon regeneration in C. elegans. Nature Communications 7, 10388 (2016).

18. Byrne, AB. et al. Inhibiting poly(ADP-ribosylation) improves axon regeneration. Elife 5, e12734 (2016).

19. Chung, S.H. et al. Novel DLK-independent neuronal regeneration in Caenorhabditis elegans shares links with activity-dependent ectopic outgrowth. Proceedings of the National Academy of Sciences 113, E2852 (2016).

20. Han, S.M., Baig, H.S. & Hammarlund, M. Mitochondria Localize to Injured Axons to Support Regeneration. Neuron 92, 1308–1323 (2016).

21. Abay, Z.C. et al. Phosphatidylserine save-me signals drive functional recovery of severed axons in Caenorhabditis elegans. Proceedings of the National Academy of Sciences 114, E10196 (2017).

22. Cartoni, R. et al. The Mammalian-Specific Protein Armcx1 Regulates Mitochondrial Transport during Axon Regeneration. Neuron 94, 689 (2017).

23. Hisamoto, N. et al. Phosphatidylserine exposure mediated by ABC transporter activates the integrin signaling pathway promoting axon regeneration. Nature communications 9, 3099–3099 (2018).

24. Kim, K.W. et al. A Neuronal piRNA Pathway Inhibits Axon Regeneration in C. elegans. Neuron 97, 511–519.e516 (2018).

25. Linton, C. et al. Disruption of RAB-5 Increases EFF-1 Fusogen Availability at the Cell Surface and Promotes the Regenerative Axonal Fusion Capacity of the Neuron. J Neurosci 39, 2823–2836 (2019).

26. Tang, N.H. et al. The mRNA Decay Factor CAR-1/LSM14 Regulates Axon Regeneration via Mitochondrial Calcium Dynamics. Curr Biol 30, 865–876.e867 (2020).

27. Yan, D., Wu, Z., Chisholm, A.D. & Jin, Y. The DLK-1 Kinase Promotes mRNA Stability and Local Translation in C. elegans Synapses and Axon Regeneration. Cell 138, 1005–1018 (2009).

28. Bounoutas, A. et al. Microtubule depolymerization in Caenorhabditis elegans touch receptor neurons reduces gene expression through a p38 MAPK pathway. Proc Natl Acad Sci U S A 108, 3982–3987 (2011).

29. Li, C. et al. The growth factor SVH-1 regulates axon regeneration in C. elegans via the JNK MAPK cascade. Nat Neurosci 15, 551–557 (2012).

30. Yan, D. & Jin, Y. Regulation of DLK-1 kinase activity by calcium-mediated dissociation from an inhibitory isoform. Neuron 76, 534–548 (2012).

31. Malinow, R.A. et al. Functional Dissection of C. elegans bZip-Protein CEBP-1 Reveals Novel Structural Motifs Required for Axon Regeneration and Nuclear Import. Front Cell Neurosci 13, 348 (2019).

32. Kaletsky, R. et al. The C. elegans adult neuronal IIS/FOXO transcriptome reveals adult phenotype regulators. Nature 529, 92–96 (2016).

33. Spencer, W.C. et al. Isolation of Specific Neurons from C. elegans Larvae for Gene Expression Profiling. PLOS ONE 9, e112102 (2014).

34. van den Brink, S.C. et al. Single-cell sequencing reveals dissociation-induced gene expression in tissue subpopulations. Nature Methods 14, 935–936 (2017).

35. O’Flanagan, C.H. et al. Dissociation of solid tumor tissues with cold active protease for single-cell RNA-seq minimizes conserved collagenase-associated stress responses. Genome Biology 20, 210 (2019).

36. Cadwell, C.R. et al. Electrophysiological, transcriptomic and morphologic profiling of single neurons using Patch-seq. Nature Biotechnology 34, 199–203 (2016).

37. Gokce, S.K. et al. A fully automated microfluidic femtosecond laser axotomy platform for nerve regeneration studies in C. elegans. PloS one 9, e113917–e113917 (2014).

38. Gokce, S.K. et al. A multi-trap microfluidic chip enabling longitudinal studies of nerve regeneration in Caenorhabditis elegans. Scientific Reports 7, 9837 (2017).

39. Wu, A.R. et al. Quantitative assessment of single-cell RNA-sequencing methods. Nature Methods 11, 41–46 (2014).

40. Ziegenhain, C. et al. Comparative Analysis of Single-Cell RNA Sequencing Methods. Molecular Cell 65, 631–643.e634 (2017).

41. Yosef, N. & Regev, A. Impulse Control: Temporal Dynamics in Gene Transcription. Cell 144, 886–896 (2011).

42. Bahrami, S. & Drabløs, F. Gene regulation in the immediate-early response process. Advances in Biological Regulation 62, 37–49 (2016).

43. Brunquell, J., Morris, S., Lu, Y., Cheng, F. & Westerheide, S.D. The genome-wide role of HSF-1 in the regulation of gene expression in Caenorhabditis elegans. BMC Genomics 17, 559 (2016).

44. Merkwirth, C. et al. Two Conserved Histone Demethylases Regulate Mitochondrial Stress-Induced Longevity. Cell 165, 1209–1223 (2016).

45. Neumann, B. & Hilliard, Massimo A. Loss of MEC-17 Leads to Microtubule Instability and Axonal Degeneration. Cell Reports 6, 93–103 (2014).

46. Savage, C. et al. Mutations in the Caenorhabditis elegans beta-tubulin gene mec-7: effects on microtubule assembly and stability and on tubulin autoregulation. J Cell Sci 107 (Pt 8), 2165–2175 (1994).

47. Duan, H. et al. Transcriptome analyses reveal molecular mechanisms underlying functional recovery after spinal cord injury. Proceedings of the National Academy of Sciences 112, 13360 (2015).

48. Teoh, J.-S., Wong, M.Y.-Y., Vijayaraghavan, T. & Neumann, B. Bridging the gap: axonal fusion drives rapid functional recovery of the nervous system. Neural Regen Res 13, 591–594 (2018).

49. Qiu, X. et al. Reversed graph embedding resolves complex single-cell trajectories. Nature methods 14, 979–982 (2017).

50. Tedeschi, A. et al. The Calcium Channel Subunit Alpha2delta2 Suppresses Axon Regeneration in the Adult CNS. Neuron 92, 419–434 (2016).

51. Tang, N.H. & Chisholm, A.D. Regulation of Microtubule Dynamics in Axon Regeneration: Insights from C. elegans. F1000Res 5, F1000 Faculty Rev-1764 (2016).

52. Chen, L. et al. Axon regeneration pathways identified by systematic genetic screening in C. elegans. Neuron 71, 1043–1057 (2011).

53. McLachlan, I.G., Beets, I., de Bono, M. & Heiman, M.G. A neuronal MAP kinase constrains growth of a Caenorhabditis elegans sensory dendrite throughout the life of the organism. PLOS Genetics 14, e1007435 (2018).

54. Xu, K., Tavernarakis, N. & Driscoll, M. Necrotic Cell Death in *C. elegans* Requires the Function of Calreticulin and Regulators of Ca^2+^ Release from the Endoplasmic Reticulum. Neuron 31, 957–971 (2001).

55. Celniker, S.E. et al. Unlocking the secrets of the genome. Nature 459, 927–930 (2009).

56. Kudron, M.M. et al. The ModERN Resource: Genome-Wide Binding Profiles for Hundreds of Drosophila and Caenorhabditis elegans Transcription Factors. Genetics 208, 937 (2018).

57. Li, C., Hisamoto, N. & Matsumoto, K. Axon Regeneration Is Regulated by Ets-C/EBP Transcription Complexes Generated by Activation of the cAMP/Ca2+ Signaling Pathways. PLoS genetics 11, e1005603–e1005603 (2015).

58. Walter, P. & Ron, D. The Unfolded Protein Response: From Stress Pathway to Homeostatic Regulation. Science 334, 1081 (2011).

59. Liu, X. et al. A Functional Non-coding RNA Is Produced from xbp-1 mRNA. Neuron 107, 854–863.e856 (2020).

60. Ritchie, F.K. et al. The Heterochronic Gene lin-14 Controls Axonal Degeneration in C. elegans Neurons. Cell Reports 20, 2955–2965 (2017).

61. Wu, Z. et al. Caenorhabditis elegans neuronal regeneration is influenced by life stage, ephrin signaling, and synaptic branching. Proceedings of the National Academy of Sciences 104, 15132 (2007).

62. Ma, T.C. & Willis, D.E. What makes a RAG regeneration associated? Front Mol Neurosci 8, 43–43 (2015).

63. Schmitt, A.B. et al. Identification of regeneration-associated genes after central and peripheral nerve injury in the adult rat. BMC Neuroscience 4, 8 (2003).

64. Chandran, V. et al. A Systems-Level Analysis of the Peripheral Nerve Intrinsic Axonal Growth Program. Neuron 89, 956–970 (2016).

65. Chen, B.K. et al. Axon regeneration through scaffold into distal spinal cord after transection. J Neurotrauma 26, 1759–1771 (2009).

66. Kadoya, K. et al. Spinal cord reconstitution with homologous neural grafts enables robust corticospinal regeneration. Nature Medicine 22, 479–487 (2016).

67. Langfelder, P. & Horvath, S. WGCNA: an R package for weighted correlation network analysis. BMC Bioinformatics 9, 559 (2008).

68. Lovatt, D. et al. Transcriptome in vivo analysis (TIVA) of spatially defined single cells in live tissue. Nature methods 11, 190–196 (2014).

69. Reynoso, M.A. et al. Translating Ribosome Affinity Purification (TRAP) followed by RNA sequencing technology (TRAP-SEQ) for quantitative assessment of plant translatomes. Methods Mol Biol 1284, 185–207 (2015).

70. Lee, J.H. et al. Fluorescent in situ sequencing (FISSEQ) of RNA for gene expression profiling in intact cells and tissues. Nature Protocols 10, 442–458 (2015).

71. Wang, X. et al. Three-dimensional intact-tissue sequencing of single-cell transcriptional states. Science 361, eaat5691 (2018).

72. Hammarlund, M., Hobert, O., Miller, D.M., 3rd & Sestan, N. The CeNGEN Project: The Complete Gene Expression Map of an Entire Nervous System. Neuron 99, 430–433 (2018).

73. Taylor, S.R. et al. Molecular topography of an entire nervous system. bioRxiv, 2020.2012.2015.422897 (2020).

74. An integrated encyclopedia of DNA elements in the human genome. Nature 489, 57–74 (2012).

75. Davis, C.A. et al. The Encyclopedia of DNA elements (ENCODE): data portal update. Nucleic Acids Res 46, D794–d801 (2018).

